# Dual-channel whole-brain imaging reveals distinct dopamine and calcium dynamics in walking *Drosophila*

**DOI:** 10.64898/2026.05.29.728103

**Authors:** Qiancheng Liang, Qiong Liu, Jing Ning, Qiantao Lv, Xingjiang Zhang, Kai Wang, Yi Sun

## Abstract

Simultaneous recording of intra- and extracellular neuronal signals across the brain during behavior is crucial for unraveling brain information processing. In *Drosophila*, large-scale recordings have been explored, yet simultaneous dual-channel whole-brain imaging remains a significant challenge. We developed a system combining a Fourier light-field microscope with dual-focal microlens arrays optimized for adult walking flies, extending imaging volume while maintaining resolution requirements. This optical system is further combined with a cross-modal 3D registration and segmentation pipeline to integrate functional and structural data. Using this system, we simultaneously measured brain-wide intracellular calcium activity and extracellular dopamine release during locomotion. We found that functional maps and neural dynamics for dopamine and calcium are distinct across the brain and within specific compartments, particularly in the mushroom body and central complex, aligning with their anatomical bases. Both calcium and dopamine representations of locomotion are distributed, yet they exhibit different patterns. Mushroom body compartments can be functionally categorized into two types based on their responses to specific locomotive actions. Forward walking or acceleration boosts activity in compartments innervated by PAM dopaminergic neurons, while dampening activity in those targeted by PPL1 dopaminergic neurons. Conversely, backward walking or deceleration heightens PPL1 activity while reducing PAM activity, consistent with the functions of these neurons in encoding approach and avoidance choices. Notably, single-compartment activity can reliably decode behavior choices. Our findings, spanning whole-brain, brain-region, and single-compartment, demonstrate the capability of our system to uncover neural dynamics across multiple scales and channels in behaving animals.

## INTRODUCTION

How different parts of the brain contribute to behavior is a fundamental question in neuroscience. Achieving this necessitates technologies capable of capturing neural activity across the entire brain in behaving animals ^1^. The adult fly brain, with its numerical tractability and genetic accessibility, is ideally suited for optically measuring whole-brain activity. Various methods have been developed for large-scale optical imaging of the fly brain. Widely used two-photon laser scanning microscopy allows for homogeneous imaging of the entire brain at high resolution ^2-6^, albeit limited by acquisition rates. Conceptually novel two-photon imaging systems such as LBM and 2pSAM offer significantly improved imaging speeds ^7,8^; however, they face limitations in axial resolution or volume size ^9,10^. Similarly, while SCAPE achieves single-objective angled light-sheet imaging ^11^, it suffers from optical aberrations that ultimately restrict the achievable imaging volume. Crucially, these microscopes rely on scanning mechanisms, and are inherently asynchronous across the imaging volume, restricting the ability to capture simultaneous neural events, which is critical for understanding circuit dynamics. Light-field microscopy captures both spatial and angular information of light rays using a microlens array ^12-14^. Its unique capability to simultaneously record the fluorescence intensity across all 3D spatial positions in a single snapshot makes it the only method capable of acquiring instantaneous 3D fluorescence intensity at the exact same moment. Light-field microscope has been used for whole-brain imaging in *Drosophila*, and has revealed complex brain-wide dynamics ^15,16^. Nevertheless, such traditional light-field microscope design exhibits reconstruction artifacts and resolution dip near the native focal plane and suffers from rapid degradation of resolution near the edges of large volumes ^12^.

Neural signals exhibit high complexity spanning molecular, circuit, and brain levels. On the molecular level, various neurotransmitters, neuromodulators, and neuropeptides permeate the extracellular space, while second messengers and other signaling molecules interact intracellularly ^17,18^. These molecular signals converge on receptors and ion channels to modulate membrane potential. Previous studies have uncovered complex relationships between intracellular and extracellular signals. For instance, while calcium is often used as a proxy for both electrical activity and neurotransmitter release ^19-21^, some studies report discrepancies between electrical activity and dopamine release ^22,23^. Deciphering this molecular complexity requires technologies for simultaneous recording across channels and space. While dual-channel imaging has been conducted in multiple regions of the fly brain ^9^, achieving it across the entire brain remains a challenge. At the circuit level, neural circuits exhibit geometric variations across animals, necessitating precise alignment to study brain-wide processing with neural circuit specificity. Furthermore, functional studies of neural circuits are often combined with structural analyses, making the identification of neural circuits from brain-wide dynamics and mapping onto anatomical atlases particularly desirable. Yet precise alignment and segmentation have been a challenge in previous whole-brain imaging studies ^9-11,15^. On the brain level, behavior and brain state are profoundly linked. Prior studies have demonstrated that neurophysiological recordings can be performed on behaving adult flies ^24,25^, facilitating the exploration of neural-behavioral relationships. So far, large-scale recordings have often been conducted either on quiescent flies ^3,9^ or have lacked dual-channel capabilities ^11,15,16^. Thus, how various signals across the brain simultaneously respond to behavioral states with neural circuit specificity remains unclear.

To address all these challenges, we designed a dual-focal Fourier light-field microscope that is fully optimized for whole-brain imaging in adult walking flies, enabling simultaneous dual-channel imaging. By performing Fourier light-field and two-photon imaging sequentially on the same animals, we implemented cross-modal registration and segmentation, allowing us to accurately analyze neural activity across the brain across channels with high precision. Using this system, we mapped dopamine and calcium activity during locomotion, revealing distinct representations across the levels of whole brain, brain regions, and individual glomeruli. Notably, we identified opposing dynamics in mushroom body compartments activated by forward and backward locomotion, corresponding to dopaminergic neurons encoding reward and punishment. Additionally, single-compartment activity derived from whole-brain imaging effectively decodes behavioral choices.

## RESULTS

### Dual-channel dual-focal Fourier light-field microscope optimized for walking adult *Drosophila* brain imaging

Whole-brain functional imaging poses challenges in the imaging volume, the spatial resolution, and the temporal resolution. Light-field microscopy achieves simultaneous 3D imaging, thereby resolving temporal-resolution limitations by imaging at a speed limited only by the camera ^12^. However, the conventional light-field microscope remains limited in its depth of imaging. Fourier light-field microscopy, by positioning the microlens array at the Fourier plane (aperture plane), achieves a spatially invariant point spread function (PSF) distribution ^26,27^. This mitigates the aliasing between angular and spatial information found in traditional light-field microscopes ^13^, thereby extending the axial imaging depth while maintaining spatial resolution stability over a broader range. Here, balancing imaging volume and the spatial resolution is a key design consideration in Fourier light-field microscopy. To mitigate imaging depth constraints imposed by the geometry of the fly brain, previous studies have often mechanically reoriented the fly head by nearly ninety degrees ^24,28,29^. While this manipulation significantly reduces the required imaging depth, the resulting unnatural posture alters sensory perception and interferes with locomotion. In contrast, in a natural walking pose, the brain extends approximately 300 μm along the optical axis. Consequently, we designed our system to accommodate this full volume, preserving the natural posture of the animal (see Methods).

By integrating a Fourier light-field with a dual-focal microlens array, we deeply optimized the system to meet the requirements for whole-brain imaging in naturally walking flies (Figure 1, Figure S1). To balance imaging volume and resolution, we optimized the effective numerical aperture of the system (see Methods). To further increase the depth of field while maintaining appropriate resolution, we used two sets of microlenses with shifted focus. To achieve dual-channel imaging, we split the fluorescence emission light after the Fourier lens and employed two sets of microlens array and camera combinations, which can be independently optimized for simultaneous recording, resulting in the <u>d</u>ual-<u>c</u>hannel dual-focal <u>F</u>ourier <u>L</u>ight-<u>F</u>ield (dcFLF) microscope.

**Figure 1.**
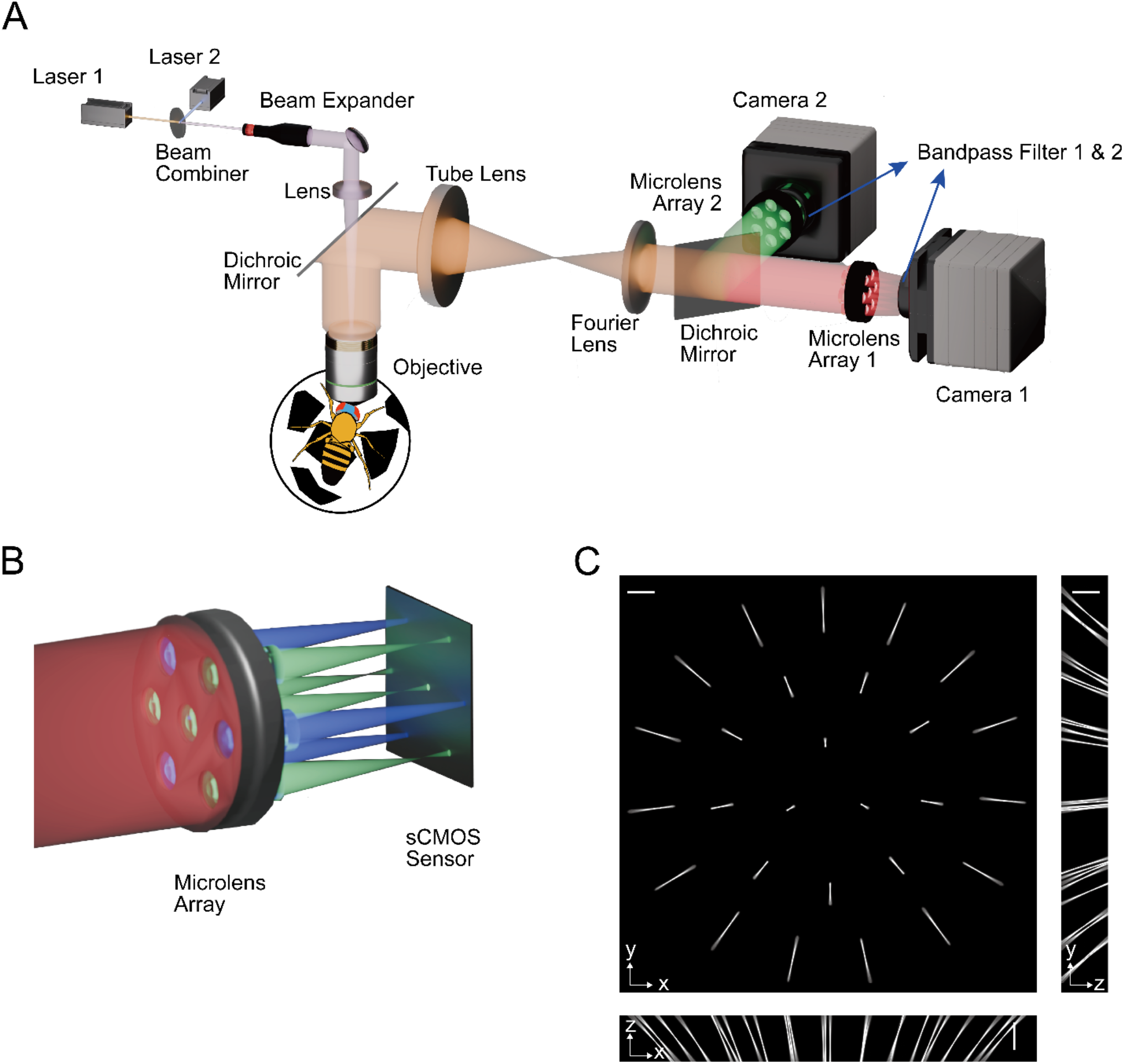
System design of dual-channel dual-focal Fourier lightfield microscope. (A) Schematic of the optical system with a fly walking on a treadmill ball. (B) Schematic of the dual-focal microlens array (MLA) in relation to the CMOS imaging sensor. The red beam on the left denotes the incident fluorescence light onto the MLA, while the blue and green beams correspond to fluorescence light focused by microlenses with two different focal lengths. Note that neither group of focused beams converges directly onto the CMOS sensor. Only seven microlenses are shown for illustrative purposes. (C) Measured PSFs of the system in the green channel, showing the maximum intensity projection of the PSF onto the x–y, y–z, and x–z planes. Scale bars are 200 µm. See Figure S1C for the PSF of the red channel.

To characterize the dcFLF system, we imaged fluorescence beads in both channels (Figure S2-3, see Methods). Using different measurement and quantification methods, we consistently found that our system achieves homogeneous resolution across a large volume. Compared to conventional light-field microscopy ^15^, our system achieves a nearly two-fold enhancement in both lateral and axial resolution (Figure S3D-E). Functional imaging speed performance is limited by the speed of indicators, which are generally below 20 Hz for widely used calcium and neurotransmitter indicators^19-21,30,31^. Thus, we imaged at 20 frames per second (fps), without temporal delay across the field. When necessary, our system can operate at over 100 fps, and is only limited by the camera. Our system enables truly simultaneous dual-channel whole-brain dynamics in adult walking flies, satisfying requirements for both imaging volume and resolution in the physiological walking pose.

### Three-dimensional cross-modal registration and segmentation

To analyze whole-brain functional imaging across channels and animals and to integrate with anatomical information requires precise registration of 3D structures. The Functional *Drosophila* Atlas (FDA) ^32^ is a reference map registered with standard anatomical maps and electron microscopy connectomes ^33-35^, providing an interface to link functional data with diverse anatomical resources. However, the FDA is based on two-photon imaging, which is challenging to register with dcFLF imaging data directly. To register the whole-brain dcFLF imaging dataset with anatomical resources, we performed cross-modal imaging with both dcFLF and two-photon imaging sequentially on the same animals (Figure 2A). To register the dcFLF volumes with the FDA, we adapted an iterative pipeline integrating both linear and nonlinear transformations and unsupervised learning ^32,36-38^ to register the imaged two-photon volumes to the two-photon-based FDA (Figure S4A, see Methods). By using two-photon imaging on the same brains as a bridge, both the dcFLF and two-photon modalities of the cross-modal dataset are registered to the FDA common space. Since the FDA is based on pan-neuronal expression of fluorescent protein via neuronal Synaptobrevin (nSyb). We used nSyb to pan-neuronally express fluorescent protein based genetically encoded indicators jRGECO1a and gDA3m ^20,31^, and achieved good registration (Figure 2A, S4B). For the newly collected dcFLF dataset, one only needs to register it to the current cross-modal dataset, without performing two-photon imaging and *de novo* registration to the FDA.

**Figure 2.**
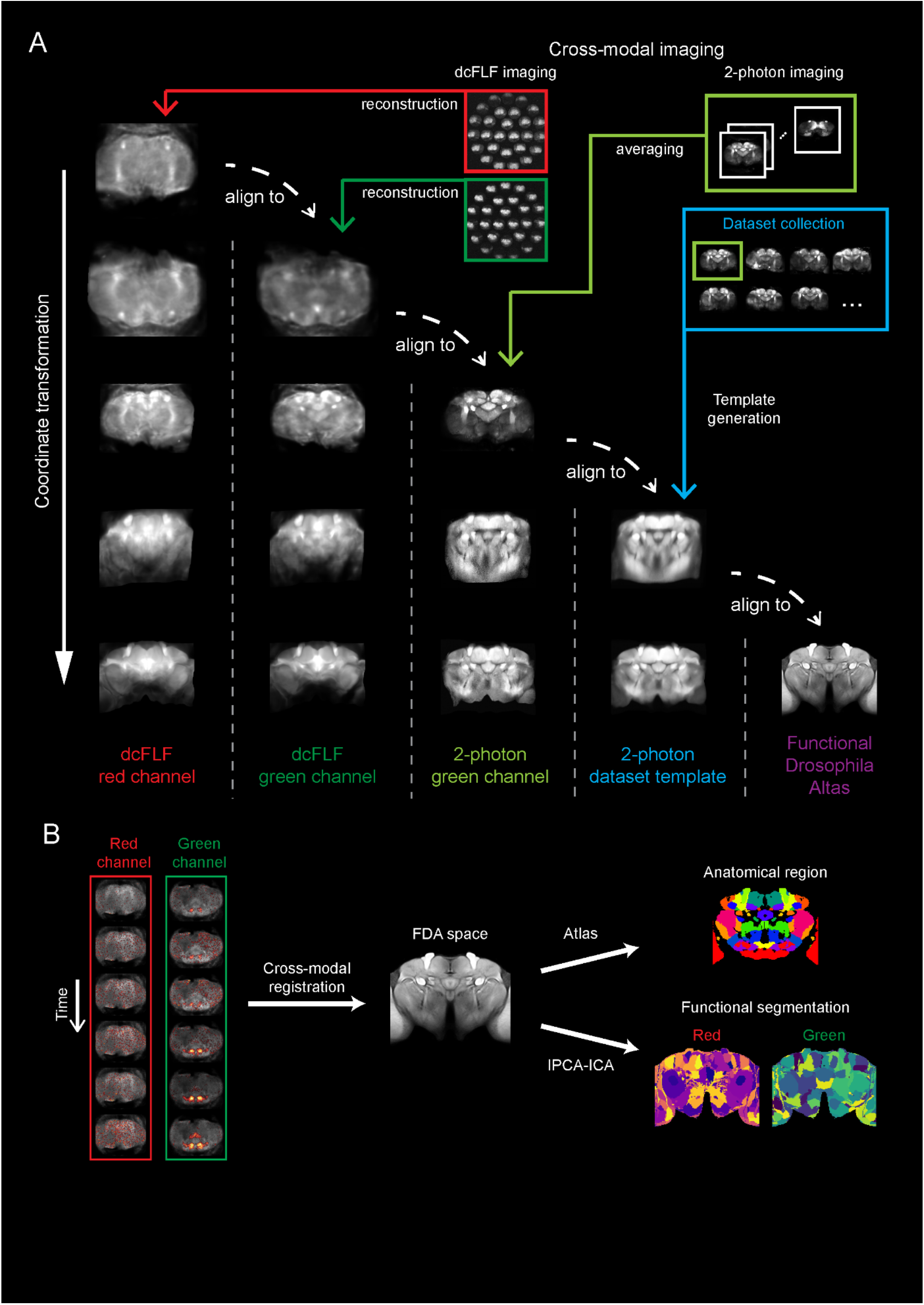
Three-dimensional cross-modal registration and segmentation pipeline for dual-channel whole-brain imaging. (A) Overview of the cross-modal registration pipeline. After dcFLF imaging and brain reconstruction, the 3D volumes in the red and green channels are first aligned. The same brain was then imaged using 3D two-photon microscopy, and the acquired two-photon volume was used as a reference to align the dcFLF volumes. By collecting a dataset of two-photon volumes from many brains and aligning all brains to a common space, a two-photon dataset template is obtained. Then, all the abovementioned volumes are aligned to the two-photon dataset template. Finally, the two-photon dataset template is aligned to the publicly available Functional Drosophila Atlas (FDA). And all the abovementioned volumes are again aligned to the FDA. (B) Overview of the pipeline for anatomical and functional segmentation. Firstly, time series dual-channel 3D volumes collected by dcFLF imaging are aligned to the FDA using cross-modal registration, as shown in A. For anatomical segmentation, an atlas containing anatomical region information is applied to the registered volumes. For functional segmentation, the IPCA-ICA approach is utilized for voxel-based segmentation in each channel.

Precise registration across multiple brains yields a significantly larger dataset, thereby enhancing the robustness of unsupervised learning for functional segmentation. To extract 3D functional structures from the whole-brain volume, we implemented a voxel-wise clustering pipeline based on neural dynamics (Figure 2B, S4C, see Methods). To alleviate memory constraints imposed by such a large dataset, we applied incremental Principal Component Analysis (IPCA) ^39^. This was followed by Independent Component Analysis (ICA) on the resulting components ^16,40^. Finally, we partitioned the brain into non-overlapping functional units by assigning each voxel to its dominant independent component. This method proved effective for segmenting functional structures in our data, as demonstrated below.

### Distinct functional clustering in calcium and dopamine systems across scales

Combining dcFLF optics with registration algorithms enables the analysis of neural dynamics across scales. By clustering neurons with correlated activity, the brain can be decomposed into distinct functional units. We next assessed the system’s capacity to resolve these structures and dynamics. To simultaneously monitor calcium and dopamine activity, we performed dcFLF imaging on flies co-expressing the green dopamine indicator gDA3m and the red calcium indicator jRGECO1a under the pan-neuronal nSyb-GAL4 driver.

The fly brain has been mapped into defined anatomical regions ^41^. To examine calcium and dopamine activity within this context, we registered our imaged volumes to the JRC2018 standard brain atlas (Figure 3A) ^34^, enabling precise anatomical segmentation (Figure 3B). Brain regions can also be defined functionally. Using the IPCA-ICA pipeline, we segmented the brain into 200 functional units for both calcium and dopamine channels independently (Figure 3C, see Methods). To validate these results, we compared functional segmentations with anatomical regions (Figure S5A). Overall, functional segmentations can be mapped onto anatomical regions in both calcium and dopamine channels (Figure S5A), validating our registration and segmentation pipeline. Since the number of functional units exceeded the number of anatomical regions, we expected finer subdivision within specific anatomical regions. Indeed, we observed dense functional parcellation concentrated in the superior neuropils, including SLP, SMP, and SIP (Figure S5A, see Table S1 for the abbreviations of brain regions). Notably, calcium and dopamine displayed distinct segmentation patterns across many of these regions (Figure 3C, Figure S5A-B).

**Figure 3.**
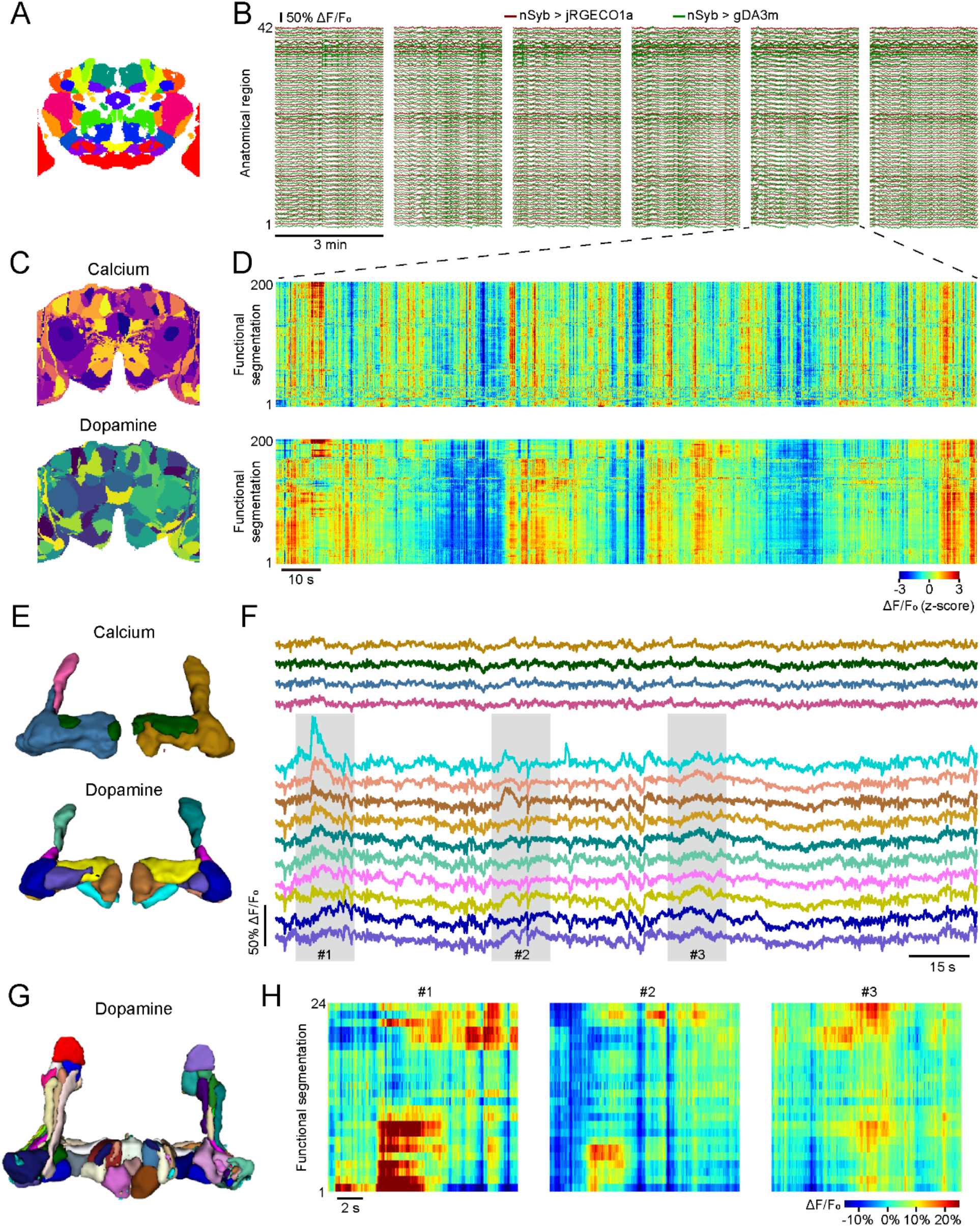
Functional clustering and neural dynamics in calcium and dopamine systems across the brain and mushroom body compartments. (A) A cross-section of the JRC2018 standard brain with anatomical segmentations. Each color denotes an anatomical region. (B) Calcium (red) and dopamine (green) response traces of the 42 anatomical regions from an example fly. Each row denotes an anatomical region, and each columnar block denotes a 3-min continuous imaging session. (C) A cross-section of the functional segmentations in calcium (upper) and dopamine (lower) channels. Same fly as in B. Each color denotes a functional segmentation. Same section plane as in (A). (D) Raster plots depicting calcium (upper) and dopamine (lower) responses of the 200 functional segmentations shown in (C), from the same period as the fifth block in (B). (E) 3D rendering of functional segmentations of the mushroom body based on brain-wide activity in calcium (upper) and dopamine (lower) channels. Same fly as in (B). (F) Calcium (upper) and dopamine (lower) response traces of mushroom body functional segmentations based on brain-wide activity as in (E), from the same period of time as in (D). Same color code as in E. Each color denotes a functional segmentation. (G) 3D rendering of functional segmentations of the mushroom body based on mushroom body-wide dopamine activity. Same fly as in (B). (H). Raster plots depicting dopamine responses of the largest 24 mushroom body functional segmentations from (G). The three blocks correspond to the same time period as the shades in (F).

To further investigate functional clustering, we zoomed in on specific brain regions. We found that distinct calcium and dopamine segmentation patterns are particularly pronounced in the dense neuropils of the central complex and mushroom body, which are higher-order regions of invertebrate brains ^42-44^. For instance, in the fan-shaped body of the central complex, functional segmentation in the calcium channel reveals a columnar pattern, while segmentation in the dopamine channel shows a tangential pattern (Figure S6B). This orthogonal layout is congruent with the columnar and tangential systems of the fan-shaped body ^45^, comprising cholinergic columnar neurons and dopaminergic tangential neurons. In the mushroom body, the calcium channel provides a rough segmentation into four regions, while the dopamine channel offers a finer segmentation with over ten regions (Figure 3E). The dopamine channel segmentation reflects the compartmentalized organization of the mushroom body, with each compartment innervated by specific dopaminergic neurons ^46,47^. We examined each segmentation individually and found a correspondence between functional segmentations and anatomically defined mushroom body compartments (Figure S6A). Given the enhanced segmentation of the mushroom body by dopamine activity, we sought to determine whether a higher resolution segmentation could be achieved. By analyzing dopamine activity specifically within the mushroom body (rather than the entire brain), we further divided it into 24 distinct segments (Figure 3G-H). These segmentations more accurately reveal the compartmentalized organization of the mushroom body (Figure S6A).

The IPCA-ICA segmentation pipeline is based on neural activity; thus, these functional segmentations provide an effective map to guide neural dynamics studies. Next, we examined calcium and dopamine dynamics across scales. Across the brain, calcium and dopamine activities exhibit distinct patterns in both anatomical and functional segmentations (Figure 3B, D). Dopamine activities appear more heterogeneous across various functional segmentations compared to calcium activities (Figure 3D). This heterogeneity is particularly evident when focusing on the mushroom body. While calcium activities are largely consistent across segmentations, we observed opposing dopamine activities in different segments (Figure 3F, H). The opposing dopamine activities may be related to the known functions of the mushroom body ^46,47^, which we will examine further below. Since calcium is present in all types of neurons – including excitatory, inhibitory, and modulatory neurons – the calcium activity measured in each voxel represents an average of these diverse and sometimes opposing activities, contributing to its apparent homogeneity.

These results demonstrate the capability of our system to reveal distinct spatial (functional clusters) and temporal (neural dynamics) structures across channels, both throughout the brain and in specific regions. Moreover, functional clustering indicates a distinct organization of spatially overlapping calcium and dopamine systems. We next applied this system to examine large-scale neural dynamics in conjunction with behavior, focusing specifically on the distinct organizations of calcium and dopamine systems.

### Distinct brain-wide calcium and dopamine maps for locomotion representation

Dopamine plays a crucial role in movement ^48,49^. In invertebrate brains, dopaminergic neurons extensively innervate the mushroom body ^50,51^. Using two-photon imaging in walking flies, calcium activity has been measured in mushroom body dopaminergic neurons ^52,53^. Additionally, direct measurements of dopamine release alongside calcium activity have been specifically investigated in the gamma lobe of the mushroom body ^54^, where dopamine release was found to be highly correlated with calcium activity in response to locomotion. Whole-brain calcium imaging reveals that movement recruits dopaminergic neurons ^6,16^. However, how locomotion affects dopamine release throughout the brain, and how dopamine release relates to calcium activity in various regions of the brain, remains poorly understood. Addressing these questions requires simultaneous measurement of whole-brain dopamine and calcium dynamics in walking flies. We set out to explore these questions using our system.

To simultaneously measure whole-brain calcium and dopamine activities, we pan-neuronally expressed jRGECO1a and gDA3m using the nSyb-GAL4 driver. We recorded locomotion velocities during imaging, confirming that transgenic flies exhibited normal walking behavior during imaging (Figure S7A-C). We measured calcium and dopamine activity for each voxel and calculated the correlation coefficient between neural activity and turning or forward walking velocities over time. By registering across brains, we obtained a brain-wide map of locomotion representation. To visualize this map, we selected four planes along the anterior-posterior axis (Figure 4A), covering many known motor regions. We plotted the locomotion correlation by voxel across the planes (Figure 4B), selecting only voxels with significant responses (P < 0.0001) for robustness.

**Figure 4.**
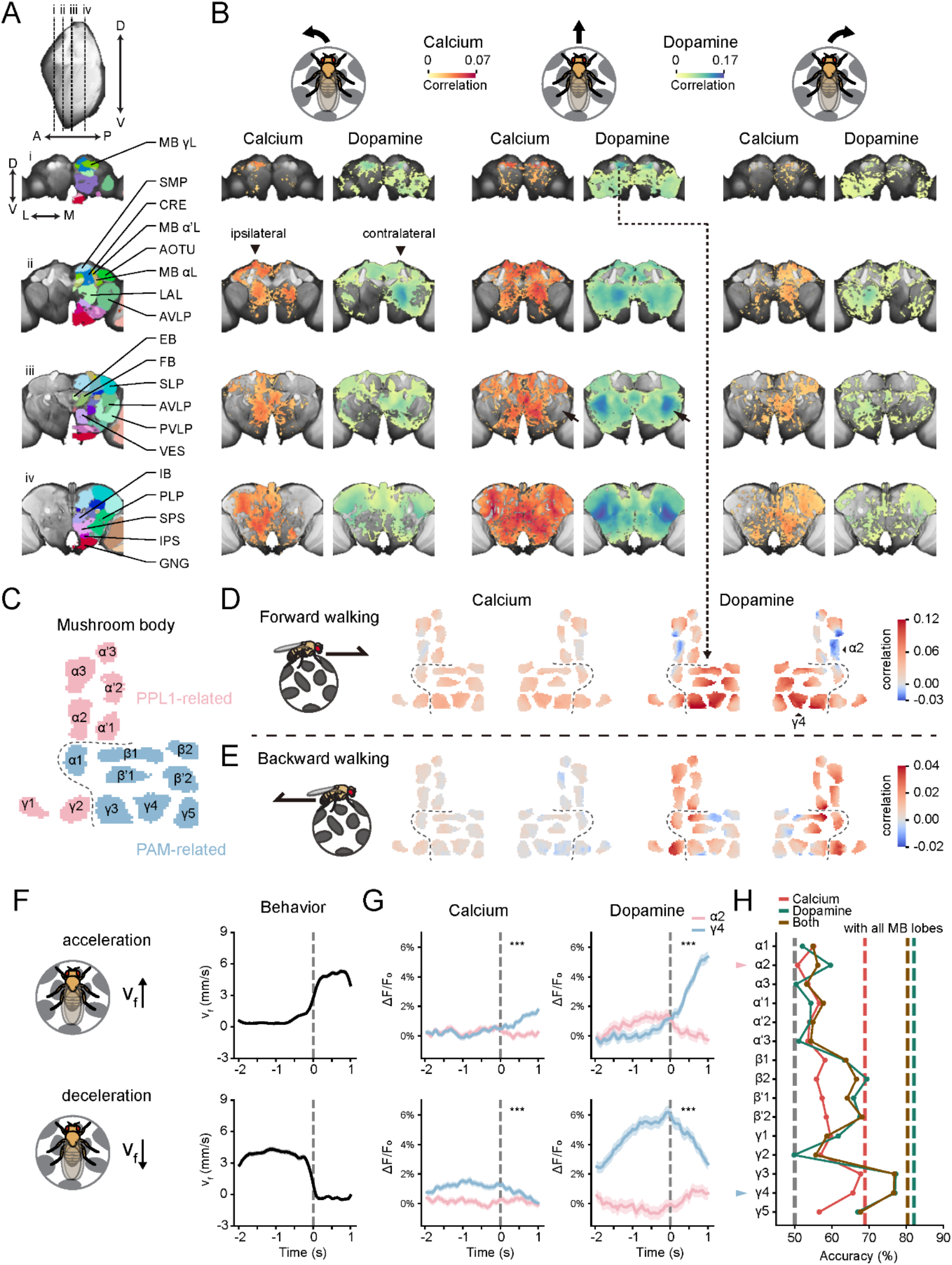
Calcium and dopamine representations of locomotion across the brain and mushroom body compartments. (A) Four frontal cross-sections of the fly brain (using FDA) from anterior to posterior (i, ii, iii, iv), with the sectioning planes marked by dashed lines on the side view of the brain at the top. Colored masks of anatomical regions are superimposed on the right half of the cross-sections, with annotations for some regions. A, anterior; P, posterior; D, dorsal; V, ventral. See Sup. Table 1 for the abbreviations of the brain regions. (B) Map of voxel-wise correlations between calcium (reddish) or dopamine (greenish) responses and turning (left and row columns) or forward (middle columns) velocities, shown as colored shades superimposed on the brain (FDA). Each row shows a cross-section corresponding to A. Only voxels with strongly significant responses (p < 0.0001, n = 12 flies) are shown. Arrowheads highlight ipsilateral calcium responses and contralateral dopamine responses to turning. Arrows highlight significant AVLP responses in dopamine but not calcium. (C) A diagram of the mushroom body compartments, with PPL and PAM innervating compartments in red and blue. (D) Map of voxel-wise correlations between forward velocity and mushroom body calcium (left) or dopamine (right) responses. Mean intensity projections along the anterior-posterior axis onto the plane in (C) by voxel are shown. (E) Same as (D), but for backward velocity. (F) Schematic and average forward velocities during acceleration and deceleration of forward velocity. Dashes denote time points with the highest velocity changes, same as follows in (G). Forward velocities shown as the mean ± S.E.M. (G) Average calcium (left) and dopamine (right) responses in mushroom body α2 (pink) and γ4 (blue) compartments during acceleration and deceleration as in F, aligned to the exact same temporal coordinates as (F). Responses shown as the mean ± S.E.M. (*** p < 0.001, Wilcoxon signed-rank test; acceleration, n = 280 trials in 12 flies; deceleration, n = 248 trials in 12 flies). (H) Classification accuracy of acceleration or deceleration, based on the activities of calcium (orange), dopamine (green), or their combinations (brown) in each mushroom body compartment using SVM. Dashes indicate the classification accuracy using data from all mushroom body compartments.

We observed distributed calcium activity across the brain during walking (Figure 4B), consistent with previous calcium imaging studies using other methods ^6,11,16^. Forward walking elicits bilaterally symmetric calcium responses, while turning triggers asymmetric calcium responses. Turning calcium responses are stronger on the side of the brain ipsilateral to the turning direction, corroborating previous reports. In the dopamine channel, we found that dopamine activity during walking is also distributed across the brain. Similar to calcium, forward walking and turning trigger bilaterally symmetric and asymmetric responses, respectively. However, unlike calcium, dopamine responses to turning appear stronger on the contralateral side (Figure 4B, arrowheads). We also observed distinct calcium and dopamine patterns during forward walking, particularly with strong dopamine responses in the posterior AVLP but not calcium (Figure 4B, arrows, correlation coefficient, dopamine 0.112, calcium 0.003). AVLP is rich in dopaminergic neurons ^55^ and is heavily innervated by ascending somatosensory inputs from the ventral nerve cord ^56,57^. Thus, AVLP dopamine activities may reflect proprioceptive feedback during movement. As discussed earlier, the responses captured by the versatile calcium reporter reflect the combined activity of various classes of neurons, which can potentially cancel each other out ^16^.

Central brain regions, such as the mushroom body and the fan-shaped body, also exhibit clear locomotion responses in both dopamine and calcium channels (Figure 4B), consistent with previous reports ^54,58^. Traditional studies of these regions relied on two-photon scanning microscopy ^53,54,59^, which is intrinsically sequential and introduces delays across the imaging field. By leveraging the temporally synchronized and spatially homogeneous imaging capabilities of our system, along with its precise registration, we next focused on neural dynamics within the mushroom body compartments.

### Opposing dopamine and calcium dynamics for locomotion representation in the mushroom body

To investigate mushroom body compartment activities during locomotion, we segmented the mushroom body into 15 compartments using a standard template (Figure 4C) ^50^. Both calcium and dopamine activities in the mushroom body are correlated with walking (Figure 4D-E). A stronger correlation was observed in dopamine activity than in calcium, consistent with reports implicating mushroom body dopaminergic neurons in movement representation ^54^. Previous brain-wide imaging studies often focused on forward walking ^6,11,16^, yet backward walking constitutes an equally critical and mechanistically distinct component of locomotion ^60,61^. Forward and backward walking can be conceptualized as approach and avoidance behaviors, potentially related to valence detection, in which the mushroom body has also been implicated ^50,53^. We observed both forward and backward walking in the flies on the treadmill during imaging (Figure S7A-C), and sought to understand how the mushroom body represents these movements. We found that the mushroom body represents forward and backward walking with distinct patterns (Figure 4D-E). Specifically, compartments in the medial horizontal lobes show a stronger positive correlation during forward walking and a weak or even negative correlation during backward walking. Conversely, opposite responses were observed in the compartments of the vertical lobes. Interestingly, these two populations correspond to the PAM and PPL1 dopaminergic neurons of the mushroom body ^62^ (Figure 4C). Consistent with this, opposing responses are more pronounced in the dopamine than the calcium channel. PAM and PPL1 neurons have been implicated in reward and punishment representation, respectively ^62^. Thus, forward walking enhances responses in PAM neurons signaling reward, while backward walking increases responses in PPL1 neurons signaling punishment. Given that forward and backward walking may indicate approach and avoidance behaviors, these observations are consistent with the known functions of PAM and PPL1 dopaminergic neurons, as well as with a recent report ^63^.

We then examined neural dynamics in individual compartments during walking, focusing on neural representations of acceleration and deceleration during forward walking. We found that both calcium and dopamine responses are compartment-specific (Figure S7D), supporting the capacity of our system to resolve compartment-specific activity. We observed opposing dynamics in both dopamine and calcium in PAM versus PPL1 populations, including in the horizontal lobe γ4 versus the vertical lobe α2 compartments (Figure 4G, S7D). These compartments are located differently across horizontal and vertical planes, requiring more time for scanning-based imaging than compartments on the same plane, such as gamma lobe neurons. Using our system, we can capture neural dynamics across the entire mushroom body at a high speed without delay, unambiguously resolving dynamics in rapid behaviors. Furthermore, because the resolution of dcFLF imaging attenuates more gradually, our system can better resolve structures located away from the geometric center of the brain, such as the mushroom body vertical lobe. Since several mushroom body compartments, such as α2, are innervated by the axons of single dopaminergic neurons, our compartment-specific results indicate single-neuron resolution.

Finally, we investigated whether the neural dynamics of an individual compartment could be used to decode locomotion decisions. We trained a support vector machine (SVM) to predict acceleration or deceleration choices based on the calcium and dopamine activity of each compartment (Figure 4H). Prediction accuracy was compartment- and channel-specific, with neural activity in most compartments exceeding chance levels. Notably, the prediction accuracy of dopamine activity tends to be higher than that of calcium and varies more across compartments. Dopamine activity in the horizontal lobe γ3 and γ4 compartments exhibits the highest single-compartment prediction accuracy, exceeding 75%. Combining data across compartments yielded higher prediction accuracy than analyzing individual compartments, whereas combining both dopamine and calcium slightly decreased prediction for both single compartments and across compartments. These results demonstrate that our compartment-resolution data can effectively predict behavior.

In summary, by resolving distinct calcium and dopamine dynamics during locomotion – from whole-brain maps to single-compartment dynamics – we illustrate how our system seamlessly scales across spatial levels and dual channels to dissect a defined behavior.

## DISCUSSION

### Simultaneous dual-channel whole-brain imaging and analysis in walking flies

Simultaneously measuring whole-brain activities across channels in behaving animals has been a long-standing goal in neuroscience ^64^. While large-scale optical imaging has been achieved in larval zebrafish, adult and larval flies, and worms ^14,65-67^, and advanced approaches have been applied to large-scale fly brain imaging ^9-11,15^ to generate valuable datasets ^2,4-6,16,29^, these methods often face trade-offs, as mentioned earlier. Optimized for normal walking posture with increased imaging depth and zero temporal delay, our dcFLF system achieves truly simultaneous whole-brain imaging in adult walking flies.

Brain function relies on the spatiotemporal interplay of neurotransmission and intracellular signaling. Capturing this complexity requires recording these processes simultaneously. While multi-channel neurochemical recording has been performed in specific circuits ^68-71^, near whole-brain dual-channel recording was only recently demonstrated in stationary flies (using calcium and acetylcholine or serotonin) ^9^, and was restricted by an imaging depth of 100 μm. Here, we present the first truly simultaneous whole-brain dual-channel recording of dopamine and calcium – a major neuromodulator and a universal second messenger. Furthermore, our system is readily adaptable to other indicators ^31,72-75^.

Comparing fine anatomical structures across channels and brains – a straightforward task when using specific and sparse drivers – poses a significant challenge in whole-brain imaging with dense pan-neuronal labeling. To resolve this, we acquired high-resolution structural scans using two-photon microscopy on the same flies imaged with dcFLF and applied a sophisticated cross-modal registration pipeline. As evidenced by mushroom body compartment level analysis, our system achieves high-precision registration of whole-brain activity across flies.

### Distinct dopamine and calcium dynamics during locomotion

Our system enabled the identification of novel functional maps and neural dynamics during locomotion at the brain, regional, and compartment levels for both dopamine and calcium. Central brain control of locomotion has been studied using large-scale calcium imaging in *Drosophila* ^6,11,16^. The role of dopamine in motor function has been investigated in both mammalian basal ganglia ^76-79^ and invertebrate mushroom body ^52,54^. However, dopaminergic neurons exhibit broad innervation across the brain in many species. Measuring brain-wide dopamine dynamics in conjunction with calcium is critical for a comprehensive understanding of motor control and the dopamine system. In fact, brain-wide dopamine release maps have been scarce across species, with previous attempts limited to tethered rats using fMRI, which suffers from restricted spatiotemporal resolution ^80^. Thus, our study provides the first brain-wide dopamine map during movement. We identified dopamine patterns that are widely distributed and unique to specific locomotive behaviors. Our findings confirmed the involvement of the mushroom body gamma lobe dopaminergic circuit in movement representation, which we extended to encompass the entire mushroom body. Additionally, we identified significant dopamine signals in other regions, including the fan-shaped body of the central complex, LAL, AVLP, and PLP. The role of dopaminergic signals in these regions in relation to motor control awaits future investigation, and our work lays the groundwork for such studies.

The mushroom body dopaminergic neurons comprise two populations: PAM and PPL1, which innervate distinct compartments and signal reward and punishment, respectively ^62^. Previous studies using two-photon calcium imaging of the gamma lobe have shown that PAM and PPL1 neurons oppositely reflect sensory stimuli and behavioral states, and oppositely regulate odor tracking ^46,53,81^. However, whether PAM and PPL1 neurons play opposing roles in locomotion representation and whether dopaminergic neurons in other lobes of the mushroom body are involved in locomotion remain unclear. We discovered that PAM neurons are activated by forward walking or acceleration, while PPL1 neurons are activated by backward walking or deceleration. Since forward walking and acceleration signal approach, while backward walking and deceleration signal avoidance, these findings potentially reflect opposing functions of PAM and PPL1 neurons in valence. Importantly, single-compartment dopamine activity, identified via the analysis of whole-brain pan-neuronal data, can be used to decode behavioral choices with high precision. Our system, equipped with compartment-resolution capabilities, allows us to dissect dopamine and calcium dynamics across the mushroom body within the broader context of whole-brain activity.

In summary, using the mushroom body as a case study, compartment-specific signals obtained from whole-brain imaging achieve a level of biologically meaningful resolution comparable to that provided by two-photon imaging focused on specific compartments. Furthermore, the same dataset can enable in-depth analysis of other regions across the brain, exploration of inter-regional interactions, and investigation of brain-wide coordination. When combined with EM connectomes using cross-modal registration, this approach will serve as a powerful tool for studying neural information processing within both local microcircuits and brain-wide networks.

### Limitations of the study

While our system successfully achieves dual-channel imaging, there is potential for further enhancement. Molecular signals occurring simultaneously are well beyond two, indicating a need to increase the number of imaging channels. Given the favorable signal-to-noise ratio in our imaging data and the availability of new spectrally variant sensors, expanding the number of imaging channels is feasible.

## Acknowledgments

We thank Y.-L. Li for sharing gDA3 fly stocks, Y.-P. Wang for technical assistance; L. Cong for assistance with the optical design; Y.-F. Pan, C. Zhou, Y. Rao for providing fly strains; B. Li, Q.-F. Ma, P.-L. Gong, L. Li, K. Si, W. Gong, S.-C. He, H.-J. Lee, J.-M. Jia, J.-Y. Peng for comments; Y.-L. Wang from Biomedical Research Core Facilities at Westlake University for support; and current and previous members of the Sun lab for discussions. This work was supported by National Natural Science Foundation of China (32371055), Research Center for Industries of the Future of Westlake University (WU2022C011), “Pioneer” and “Leading Goose” R&D Program of Zhejiang (2024SSYS0031), Key Laboratory of Growth Regulation and Translational Research of Zhejiang Province, to Y.S.; and National Natural Science Foundation of China (32125020) to K.W.

## Author contributions

Conceptualization, Y.S., K.W.; methodology, Q.-C.L., Q.L., J.N., X.-J.Z., K.W., Y.S.; investigation, Q.-C.L., Q.L., J.N., Q.-T.L., X.-J.Z., Y.S.; data curation, Q.L., J.N., Q.-C.L.; formal analysis, J.N., Q.L., Q.-C.L., Y.S.; visualization, Q.-C.L., Q.L., J.N., Y.S.;writing-original draft, Y.S., Q.L., Q.-C.L., J.N., K.W.; project administration, Y.S., Q.-C.L., Q.L.; funding acquisition, Y.S., K.W.; resources, Y.S.; supervision, Y.S.

## Declaration of interests

The authors declare no competing interests.

## METHODS

### EXPERIMENTAL MODEL AND STUDY PARTICIPANT DETAILS

#### Fly stocks

All imaging experiments were performed on virgin female fruit flies (*Drosophila melanogaster*). Flies were reared on standard cornmeal food in 35 mL vials, single-housed, aged 4-6 days after eclosion, and raised on a 12:12 dark: light cycle at 25 ℃ and 60% humidity. For pan-neuronal expression of jRGECO1a and gDA3m, flies of the genotype were used: w+; UAS-jRGECO1a/+; nSyb-GAL4/UAS-gDA3m.

## METHODS DETAILS

### dcFLF microscope optical system design

#### Optimizing Effective Numerical Aperture of the Optical System

To balance the requirements between imaging volume and spatial resolution, we optimized the effective numerical aperture of the system (*NA*_*sys*_), based on the fundamental trade-off between depth of field (*DOF*) and lateral resolution *R*_*xy*_. The *DOF* is defined as ^82^:

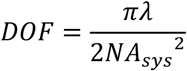

where *λ* denotes the wavelength of a specific fluorescence channel. Meanwhile, the lateral resolution *R*_*xy*_ is governed by ^82^:

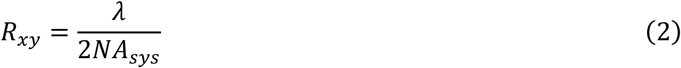

It can be seen from formulas (1) and (2) that DOF and lateral resolution are antagonistic.

To evaluate whether the selected *NA*_*sys*_ meets the spatial resolution and DOF requirements for fly whole-brain imaging, we performed a numerical simulation of diffraction using the Angular Spectrum Method (ASM) ^83^. ASM provides a more rigorous solution by operating in the spatial frequency domain. It decomposes the initial light field into a superposition of plane waves, and the propagation process is reduced to a phase-shift operation in the Fourier domain:

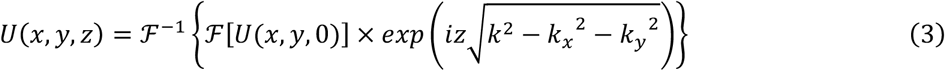

Here, *U(x, y, 0)* and *U(x, y, z)* denote the complex amplitudes of the optical field at the initial plane (z = 0) and the observation plane after propagating a distance *z*, respectively. The wave number *k* is defined as 2*π*/*λ*, where *λ* denotes the wavelength of the fluorescent channel. The terms *k*_*x*_ and *k*_*y*_ denote the transverse spatial angular frequencies, corresponding to the directional components of the propagating plane waves in the Fourier domain. The exponential term 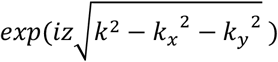 serves as the free-space transfer function, in which the radical term characterizes the longitudinal wave vector component 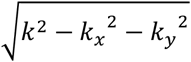, accounting for the phase accumulation during propagation. By applying an inverse Fourier transform, the propagated optical field is recovered in the spatial domain. Given that fluorescence microscopy operates as an incoherent imaging system, the detector captures the optical intensity rather than the field amplitude. Therefore, the Point Spread Function (PSF) is manifested as the squared modulus of the complex amplitude *U(x, y, z)*:

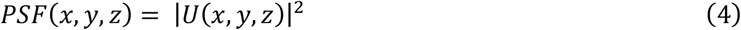

Simulating PSFs under diverse *NA*_*sys*_ configurations allowed us to validate the system resolution limits at various axial depths (*z*), ensuring compliance with fly whole-brain imaging requirements during the initial design phase.

#### Optical design via resolution and aperture matching

We designed the tube lens and Fourier lens by ensuring resolution matching throughout the system. We first selected the objective and the camera. To mitigate inherent signal attenuation and the fill-factor limits of the microlens array (MLA) mechanical housing, we optimized the detection path using a high numerical aperture (*NA*_*obj*_) objective and a high quantum efficiency sCMOS camera. We then determined the optimal focal lengths of the tube lens and Fourier lens based on a dual-matching approach.

1. **Resolution matching**: To satisfy the Nyquist-Shannon sampling criterion, we matched the optical resolution to the pixel size of the sCMOS sensor. This ensures the full utilization of both optical resolution and camera sampling resolution.
2. **Aperture matching**: We optimized the relay optics to scale the objective rear pupil to the active area of the sCMOS sensor. This matching maximized the effective use of both the optical system and the camera sensor.

#### Dual-focal microlens array design

We derived the diameter and initial focal length of the microlens from the system magnification *M* and the system effective numerical aperture *NA*_*sys*_. To optimize the MLA geometry, we compared rectangular, hexagonal, and concentric circular arrangements. We simulated the PSFs of optical systems using these MLA designs via ASM diffraction modeling, as discussed above, and performed 3D reconstructions. Based on the spatial resolution of the simulated results, we chose the concentric circular layout. We further employed this simulation-reconstruction procedure to fine-tune the focal lengths of the two microlens groups of the dual-focal microlens array. The dual-focal microlens design is illustrated in Figure 1B, and the layout of the microlens array is shown in Figure S1A.

### dcFLF microscope optical system implementation

In the fluorescence excitation path, two continuous-wave lasers (473 nm and 561 nm; Laser Quantum, Stockport) were coaxially combined using a dichroic beam combiner (86-392, Edmund Optics, Barrington). The beam diameter was expanded via a beam expander (GBE15-A, Thorlabs, Newton) to ensure a uniform excitation field across the sample. This beam was relayed through a 4f system consisting of an achromatic lens (f = 125 mm; 49-361, Edmund Optics) and an objective (25×/NA 1.1, CFI75 Apo 25 XC W, Nikon, Tokyo). A dichroic mirror (Di01-R488/561-25×36, Semrock, Rochester) redirected the excitation light into the objective while transmitting the emitted fluorescence. Excitation power was modulated using a continuously variable neutral density filter (GCO-0703M, DHC, Beijing). In the fluorescence detection path, the fluorescence light was collected by the objective and redirected horizontally via a dielectric mirror (39-226, Edmund Optics). We selected a high quantum efficiency sCMOS camera (Dhyana 400BSI V2, Tucsen, Fuzhou). Another 4f relay system (AC508-200-A, Thorlabs; 49-361, Edmund Optics) was employed for resolution and aperture matching (see above). A custom-designed microlens array (MLA, see above) was placed at the conjugate plane of the objective rear pupil. The schematic, optical path, and component diagrams of the system are shown in Figure 1A, Figure S1A, and Figure S1B, respectively.

### dcFLF microscope spatial resolution and PSF characterization

To validate system performance for 3D neural activity mapping throughout the *Drosophila* brain, we conducted a multidimensional characterization of spatial resolution. Given that dcFLF exhibits depth-dependent imaging properties, characterizations were performed at multiple axial depths.

#### System spatial cutoff frequency

In incoherent imaging systems, such as fluorescence microscopy, the Optical Transfer Function (OTF) is calculated as the Fourier transform of the PSF:

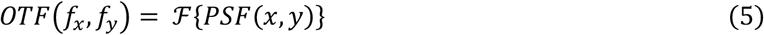

The spatial cutoff frequency (*f*_*c*_) represents the threshold at which the OTF magnitude drops to zero. Beyond this threshold, the system can no longer modulate spatial information, resulting in a complete loss of image contrast for features above this frequency bound. To determine whether the system maintains a consistent spatial cutoff frequency throughout the designated 3D imaging space, we characterized the system’s PSF using fluorescent beads at multiple locations. We systematically sampled point-source responses at defined spatial intervals across the FOV (x = 0, ±50, ±100, ±150 μm) and depth (z = 0, ±50, ±100 μm). This localized PSF analysis enables verification of spectral coverage stability within the specified imaging depth and range. The frequency-domain representation (OTF magnitude) was obtained by applying a two-dimensional Fast Fourier Transform (2D-FFT) to the measured PSFs. The system’s effective cutoff spatial frequency was estimated by measuring the diameter (*D*_*OTF*_) of the high-frequency response region in the OTF map:

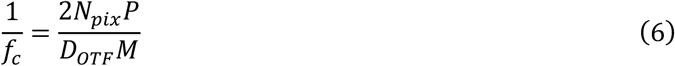

where *N*_*pix*_ is the number of pixels (sampling points in the frequency domain), *P* is the physical pixel size of the sCMOS sensor, and *M* is the system magnification. The effective response regions of the OTF at different locations of the volume were identified in Figure S2A-B, demonstrating consistency.

#### Characterizing axial resolution using computational superposition

While spatial resolution is conceptually defined as the minimum distance required to resolve two point sources along a specific axis, the precise axial positioning of fluorescent beads with micrometer-scale accuracy remains technically demanding ^26^. To circumvent this, we employed a computational superposition strategy to characterize the axial resolution of our system. Specifically, we acquired image stacks of a planar distribution of fluorescent beads at discrete axial depths. By superimposing pairs of these images, we generated synthetic 3D volumes that effectively simulate two layers of beads with precisely controlled axial displacements around a target depth. We then performed 3D reconstructions on the superimposed datasets to define the axial resolution limits for each layer based on a computationally displaced pair of beads. As shown in Figure S2C– D, these displacements enable clear separation of the green and red channels into two distinct entities. The results (Figure S2E) demonstrate that the system can resolve axial distances down to 4 μm in both the green and red channels.

#### Volumetric resolution characterization via FWHM analysis

Beyond the classical criterion of distinguishing adjacent point sources, we quantified the spatial resolution by measuring the Full Width at Half Maximum (FWHM) of the reconstructed intensity distribution from fluorescent beads. This metric provides a robust simultaneous assessment of both lateral and axial imaging performance (Figure S3A). To map the resolution profile throughout the entire imaging volume, we positioned a fluorescent bead at multiple spatial locations within the imaging volume. Following 3D reconstruction, we calculated the FWHM of the beads at these discrete locations to characterize the local spatial resolution in two channels (Figure S3B-C).

#### Resolution characterization using reconstruction of multiple microspheres

Instead of positioning a specific fluorescent bead at particular coordinate, we used agarose (1%) to immobilize multiple stochastically distributed 2 μm fluorescent beads to characterize 3D resolution at different depths. We reconstructed 3D images of these beads, and determined the lateral and axial resolutions at different depth based on the FWHM of the images. Due to the stochastic spatial distribution of the beads, we sampled all discrete beads across the axial range, followed by non-parametric linear regression and then Lowess filtering with a span of 60% to obtain the resolution characterization (Figure S3D-E).

#### Measurement of the Point Spread Function

To account for system-specific aberrations, PSFs were experimentally determined. Unlike conventional light-field microscopes, Fourier light-field microscope exhibits spatial invariance, enabling the measurement matrix to be represented by a single PSF, simplifying PSF measurement. Given the theoretical resolution of the system, we measured PSFs using 2 μm fluorescent beads. PSFs were acquired over a 300 μm axial range with a 2 μm step size. During calibration, the symmetry of the axial maximum intensity projections (MIPs) was monitored to ensure precise alignment of the MLA with the optical axis.

### dcFLF microscope 3D image reconstruction

3D volumes were reconstructed using a Richardson-Lucy deconvolution algorithm ^26^. The reconstruction algorithm implemented forward and back-projections in the frequency domain via FFT. For iteration *i+1*^*th*^, the estimate of the 3D volume in *k*^*th*^ layer (where *k* ∈ {1 … … *n*}), *g*_*i*+1_(*x*, y, z_*i*_) was updated according to the following iterative scheme:

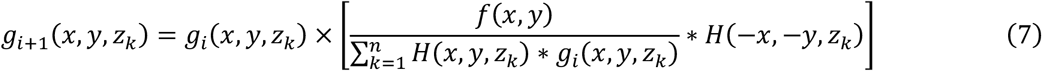

where *f(x,y)* denotes the measured 2D light-field image, *H(x,y,z*_*k*_) denotes the forward-projection operator (the measured PSF). Prior to deconvolution, raw images underwent preprocessing, including background subtraction.

To prevent noise amplification while maintaining structural integration, the deconvolution process was typically terminated after a specific number of iterations. All convolution operations were computed in the frequency domain using FFTs for efficiency. The reconstruction algorithm was implemented in MATLAB and accelerated using GPU (GeForce RTX 3080, NVIDIA).

### Fly surgery

A custom-made holder plate was constructed from an etched stainless-steel foil (0.03 mm thick) with a 0.8 mm-diameter central aperture, folded into a pyramidal geometry, and bonded to an aluminum base. We also custom-designed, fabricated, and assembled a copper surgery station that includes a crossbar with a groove to trap the fly’s body. The surgery station was maintained at 2 ℃ using thermoelectric coolers with a PID controller. After brief anesthetization on ice, flies were positioned on the surgery station and tethered to the holder plate using UV-curable glue (Norland Optical Adhesive 68). To reduce brain movements, we glued the proboscis using UV-curable glue. Flies were then allowed to recover from anesthesia for 30-60 min before surgery. The holder plate was filled with extracellular saline (103 mM NaCl, 3 mM KCl, 1 mM NaH_2_PO_4_, 5 mM TES, 26 mM NaHCO_3_, 4 mM MgCl_2_, 2.5 mM CaCl_2_, 10 mM trehalose, and 10 mM glucose, pH 7.4 with osmolarity adjusted to 270–275 mOsm). The head cuticle was cut open with a sharp needle and removed with forceps. The trachea and fat tissues were removed to expose the central brain. Flies were allowed to adapt to walking on a treadmill ball for 10-20 min before imaging experiments.

### dcFLF functional imaging

To synchronize imaging with auditory stimulation and locomotion recordings, the sCMOS camera was externally triggered via a data-acquisition card (PCIe-6321, National Instruments) using custom scripts in MATLAB. To ensure proper writing of buffered images to the hard drive, we recorded in blocks of approximately two hundred seconds each. For each fly, a total of six blocks were recorded. For functional imaging, we recorded at 20 frames per second (fps) with an exposure time of 48 ms. To position the brain at the center of the imaging volume while ensuring the fly walks reliably on the treadmill ball, we adjusted the fly’s position with respect to both the objective and the ball. We captured sample images, enabling us to quickly reconstruct the brain’s 3D structure via remote access to the GPU workstation. We used the easily identifiable structure of the mushroom body as the landmark for brain positioning.

### Two-photon structural imaging

Flies were immediately transferred to perform two-photon structural imaging after dcFLF imaging. We collected imaging data with a custom-built two-photon microscope with a water-immersion objective (25x, NA 1.1, Nikon) and ThorImage (Thorlabs) for image acquisition. An ultrafast laser (InSight X3, Spectra-Physics) was used to excite jRGECO1a and gDA3m simultaneously at 1000 nm. The fluorescence emission signals were collected and amplified by a pair of photomultiplier tubes (PMTs) through a 562 nm long pass dichroic and a 525/50 nm bandpass filter (green channel) or a 607/70 nm bandpass filter (red channel). 100 imaging volumes (341.21 x 341.21 x 301 μm) were acquired at a resolution of 512 x 512 x 301 voxels.

### 3D registration pipeline

#### Within-animal registration

##### dcFLF structural brain generation and motion correction

For cross-channel alignment and motion correction, a structural brain was first constructed for each channel of each fly. To achieve this, we reconstructed the volumes and selected volumes at 1000-sample intervals from the time series of functional volumes. These selected volumes were averaged, followed by warping each volume to the average using linear (affine) and non-linear (SyN) transformations in ANTs. Then, the warped volumes were averaged again, and each selected volume was realigned to this new target. Volumes after this second registration were averaged again, and the resulting averaged volume was used as the dcFLF structural brain. For motion correction, the time series of functional volumes in each channel were registered to their corresponding dcFLF structural brain through an affine transformation using ANTs.

##### Cross-channel alignment

To align the green and red channels of dcFLF imaging, we selected the green channel (gDA3m) as the registration target because its clearer two-photon structural brain facilitates subsequent cross-modal alignment. We first manually aligned the dcFLF structural brain in the red channel to that in the green channel using rigid transformations (translation, rotation, and flipping) in ITK-SNAP (www.itksnap.org) ^84^, and obtained the transformation parameters. Using these parameters as the initial transformation, we then performed SyN registration to align the red channel to the green channel in ANTs, obtaining the final parameters. The motion-corrected functional volumes of the red channel were registered to the green dcFLF structural brain with these final parameters in ANTs.

##### Two-photon structural brain generation

Following a procedure similar to that for dcFLF structural brain generation, 100 structural volumes from the green channel were motion-corrected and averaged to create a two-photon structural brain, via one round of alignment with affine and SyN transformation in ANTs.

##### Cross-modal alignment

To register the green channel across the dcFLF and two-photon imaging modalities for each fly, we first performed a coarse alignment of the dcFLF structural brain to the two-photon structural brain using the landmark registration tool in 3D Slicer (www.slicer.org) ^85^. By manually marking specific landmarks of the mushroom body and central complex, we derived the initial transformation parameters. These parameters were then used to initialize a fine registration between the dcFLF structural brain and the two-photon structural brain through SyN transformation using ANTs. The resulting transformation parameters were applied to the functional volumes of both channels in green dcFLF structural brain coordinate, thereby uniformly transforming all functional imaging data into the coordinate of the two-photon structural brain.

#### Cross-animal registration

To align brain volumes from all flies to the Functional Drosophila Atlas (FDA), we employed a two-step registration process consisting of coarse and fine alignment. Since our two-photon imaging was conducted in a posture different from that of the FDA, we first generated a dataset template and then accurately registered it to the FDA. To accommodate the two-photon imaging field of view, we adjusted the FDA by cropping most of the optical lobes. To improve computational efficiency, we downsampled the FDA to 456 x 384 x 240 voxels. The coarse alignment involved a rough registration of each animal’s two-photon structural brain to the adjusted FDA using the landmark registration tool in 3D Slicer. This involved manually marking specific landmarks of the mushroom body and the central complex. The resulting transformation parameters were then applied to all functional volumes in the coordinate of the two-photon structural brain, transforming them into an intermediate coarse-aligned coordinate. Subsequently, a dataset template was generated from the coarse-aligned two-photon structural brains of all flies in the dataset and registered to the adjusted FDA via a combination of linear (affine) and non-linear (SyN) transformations, along with unsupervised learning (SynMorph), using BIFROST ^32,36-38^. Finally, the transformation parameters calculated for each animal during this process were applied to the functional volumes in the intermediate coordinate for the fine alignment. In this way, the dcFLF functional imaging data from both channels for all flies were registered to the FDA coordinate system. To reduce storage occupancy, the final aligned volumes were downsampled to 152 x 128 x 80, matching the initial resolution of the dcFLF-reconstructed volume. For parallel processing, we implemented the registration pipeline using the Slurm cluster scheduling system.

#### Aligning anatomical regions and MB compartments

We obtained anatomical region labels in the adjusted FDA coordinates based on the neuropil mask in the JRC2018Unisex coordinate ^34^ from the Virtual Fly Brain (v2.virtualflybrain.org). We first aligned the neuropil mask to JRC2018F in ANTs using the JRC2018Unisex-to-JRC2018F transformation from Janelia (https://www.janelia.org/open-science/jrc-2018-brain-templates). Subsequently, we registered the JRC2018F template brain to the FDA using BIFROST, thereby obtaining the corresponding transformation parameters. The neuropil mask in the JRC2018F coordinate was then transformed to the FDA space by BIFROST using the same parameters with nearest-neighbor interpolation. Finally, the wrapped mask was adjusted to the imaging volume and downsampled to improve computational efficiency, as mentioned above from the FDA to the adjusted FDA.

To obtain the MB compartment labels in the adjusted FDA coordinates, we used the MB compartment mask ^50^ and a procedure analogous to that applied for the anatomical regions mentioned above. Since the MB compartment mask was in JFRC2013 coordinates, we registered JFRC2013 to JRC2018F in the first step.

#### 3D registration evaluation

To evaluate registration accuracy, we assessed the brain-to-brain variation using 6 manually labeled landmarks in 3D space, including the right and left BU, the center of the bilateral mushroom body horizontal lobes, the center of EB, and the top of the right and left MB vertical lobes (Figure S4B). For each fly, we labeled these landmarks (using a round brush with an 8-voxel size) in the dcFLF structural brain of the green channel using ITK-SNAP, then transformed this label mask to the adjusted FDA using the corresponding parameters from the 3D registration pipeline. The centroid of the labeled region post-registration served as the position of each structural landmark, and we calculated pairwise Euclidean distances across all flies for each structure.

### Functional segmentation and analysis

#### Functional segmentation

To identify the boundary of the brain, we selected voxels with intensity values above 0.1 from FDA to construct a brain mask. To focus on central brain regions, we excluded the optic lobes from this mask, resulting in a final voxel set containing over half a million valid voxels. We applied this mask to each imaging volume to extract voxel-level time series of activity. To correct for photobleaching, we estimated the baseline fluorescence F_0_ for each time series by applying a 1-minute moving average filter, and subtracted the baseline from the time series. We then computed ΔF/F as (F − F_0_)/F_0_. Voxels exhibiting abnormally high signal fluctuations were excluded from further analysis. We augmented the dataset by leveraging the brain’s bilateral symmetry, mirroring the data along the midline to effectively double its size. We then employed an IPCA/ICA pipeline ^16,39,40^, implemented with Python on a workstation. Given the large scale of the dataset— approximately 300 billion voxels in our dataset —we performed incremental PCA and retained the first 200 principal components. This number was chosen as a conservative threshold (approximately twice the elbow point in the log singular-value spectrum, which is around 100) to preserve activity-related signals. Subsequent ICA decomposition of the principal component maps yielded 200 independent components. Finally, we assigned each voxel a cluster label corresponding to the independent component with the highest loading weight. Each cluster represents a distinct functional segmentation associated with a unique, coherent spatiotemporal activity pattern.

#### Brain-wide comparison of functional segmentations and anatomical regions

To assess the correspondence between functional segmentations and anatomical regions, we calculated the voxel percentage of each anatomical region within every functional segment, excluding unlabeled voxels (Figure S5A). A segment was assigned to a specific anatomical region if that region accounted for more than 40% of its total voxels. To visualize this relationship, we projected the segment along the anteroposterior (AP) axis and overlaid it onto a partially exploded brain map, preserving the spatial position of each functional segment relative to its dominant anatomical region (Figure S5B). To remove tiny fragments while retaining prominent structures, we performed morphological filtering using a closing operation (2 iterations, square connectivity = 1) followed by an opening operation (2 iterations, square connectivity = 1).

#### Visualization of the functional segmentation of MB

To visualize MB functional segmentations, we rendered segmentations derived from clustering either whole-brain calcium or dopamine activity (Figure 3E), or exclusively MB dopamine activity (Figure 3G). To establish the correspondence between functional segmentations and anatomical compartments in MB, we calculated the structural similarity between each functional segment (split into two unilateral parts when spanning both hemispheres) and various combinations of anatomical compartments, and presented the combinations yielding the highest similarity (Figure S6A). The structural similarity between two voxel sets was quantified by the Sørenson–Dice coefficient, defined as:

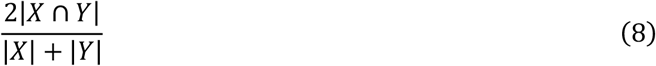

Where |*X* ∩ *Y*| denotes the intersection size between the two sets, while |*X*| and |*Y*| denote their respective cardinalities.

### Locomotion monitoring and processing

We recorded the locomotion behavior of the fly during dcFLF imaging by measuring the rotations of an air-supported polyurethane ball (8 mm diameter) on which the fly walked. The ball rested on a custom-made aluminum holder with a semi-spherical concave, at the bottom of which a 1 mm channel was drilled to supply air flowing at 0.5 L/min, controlled by a glass rotor flowmeter (LZB-3WB, Kede Instruments) and mass flow controller (D07-7, Beijing Sevenstar Flow). The position of the treadmill ball was adjusted for each fly guided by a camera (HBVCAM-W202012HD V33, HUIBER VISION TECHNOLOGY) placed on the side relative to the objective. To promote walking during the experiments, we heated the aluminum holder and delivered sound stimulation to the fly. The holder was heated to maintain an air temperature of approximately 29 ℃ at the apex of the treadmill ball, where the fly was positioned ^6,16^. We delivered sound via two speakers at the back. The sounds included pulse, sine, and white noise, each lasting 1 s, along with their combinations, presented in a random order. Sound stimuli were generated at a sampling frequency of 10 kHz using custom MATLAB code. To ensure that the antennae and aristae were mobile and thus sensitive to auditory stimulation, we checked each fly with gentle airflow. To monitor locomotion, we illuminated the fly and the ball with a ring-shaped infrared LED light source and captured videos from the front side with an industrial camera (MV-CA013-21UM, Hikrobot) at 50 fps using a zoom lens (MVL-HF3528M-6MP, Hikrobot). To synchronize locomotion recording with functional imaging and sound stimulation, we externally triggered the industrial camera, as mentioned above. We marked the treadmill ball with random patterns, and used Fictrac 2.0 ^86^ to track ball rotation in real time. In cases where the tracked data contained errors, the original recordings were reprocessed offline. For time intervals in which errors remained despite multiple re-tracking attempts, the corresponding velocity values were assigned null values (constituting less than 0.1% of the total dataset) and excluded from further analyses. The velocity was smoothed using a Savitzky–Golay filter with a window length of 500 ms and a third-order polynomial.

### Analysis of neural activity and locomotive behavior

#### Voxel-wise correlation analysis

To examine the relationship between neural activity and locomotion, we calculated voxel-wise Pearson correlation coefficients between calcium or dopamine signals and four velocity components: forward, backward, left, and right. Velocity was decomposed into these non-negative directional components and downsampled to align with the imaging frame rate (from 50 to 20 Hz). Correlations were computed using a 100-frame (5 s) rolling window and subsequently averaged across all windows for each animal. For each voxel, we calculated the mean value across all animals.

#### Forward and backward walking bout detection

To detect bouts of forward and backward walking (Figure 4D, Figure S7B-C), we calculated the ratio of forward to backward walking using a 3-s sliding window (spanning 2 s prior to 1 s following each sampling moment) with a step size of 5 frames (100 ms):

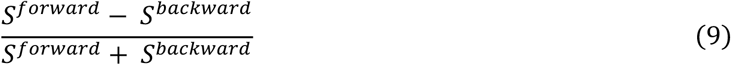

where *S*^*forward*^ and *S*^*backward*^ denote magnitudes of forward and backward displacement within this time epoch, respectively. Forward walking bouts were identified as ratios above 0.5, and backward walking bouts as ratios below -0.5. The average of the first 2.5 s across all bouts was used to illustrate sustained forward and backward walking (Figure S7B-C).

#### Acceleration and deceleration bout detection

To detect bouts of acceleration and deceleration in forward velocity (Figure 4F-H), we calculated a normalized forward velocity difference before and after the sampling moment using a 3-s sliding window (spanning 2 s prior to 1 s following each sampling moment) with a step size of 5 frames (100 ms):

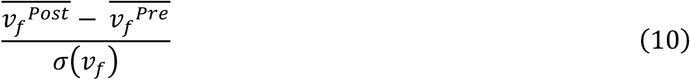

where 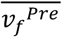 and 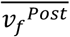 denote the mean forward velocities during the 2-s interval preceding and the 1-s interval following the sampling moment, respectively, and *σ*(*v*_1_) denotes the standard deviation of forward velocity within the corresponding consecutive recording block. Bouts of acceleration or deceleration were identified by detecting peaks (absolute amplitude > 2) in this normalized forward velocity difference. The average of the corresponding calcium and dopamine activities during these 3-s bouts was shown (Figure 4G and S7D). To quantify the difference in α2 and γ4 response changes during acceleration and deceleration (Figure 4G), we separately calculated the difference in average activity of α2 and γ4 before and after the sampling moment.

#### Decoding behavioral choices using MB compartment activity

To classify forward velocity changes of acceleration or deceleration using MB calcium or dopamine signals, we employed a Support Vector Machine (SVM) classifier. For each channel and each compartment, we first used PCA to reduce the dimensionality of the signal change during the bout, which was a time series of 60 data points. We used the first 10 principal components (PCs) as input features for an SVM classifier to predict bout changes. The accuracy was determined via 5-fold cross-validation. We further assessed the accuracy of two combination strategies: combining activity across all MB compartments and concatenating data from the two channels (Figure 4H).

## QUANTIFICATION AND STATISTICAL ANALYSIS

We used non-parametric tests for statistical analysis. To compare two groups, we used the Wilcoxon signed-rank test. To assess whether observed values were statistically different from zero, the Wilcoxon one-sample signed-rank test was used. All statistical analyses were performed in Python.

## Notes

### Competing Interest Statement

The authors have declared no competing interest.

