## Supplemental Information for "Dual-channel whole-brain imaging reveals distinct dopamine and calcium dynamics in walking *Drosophila*"

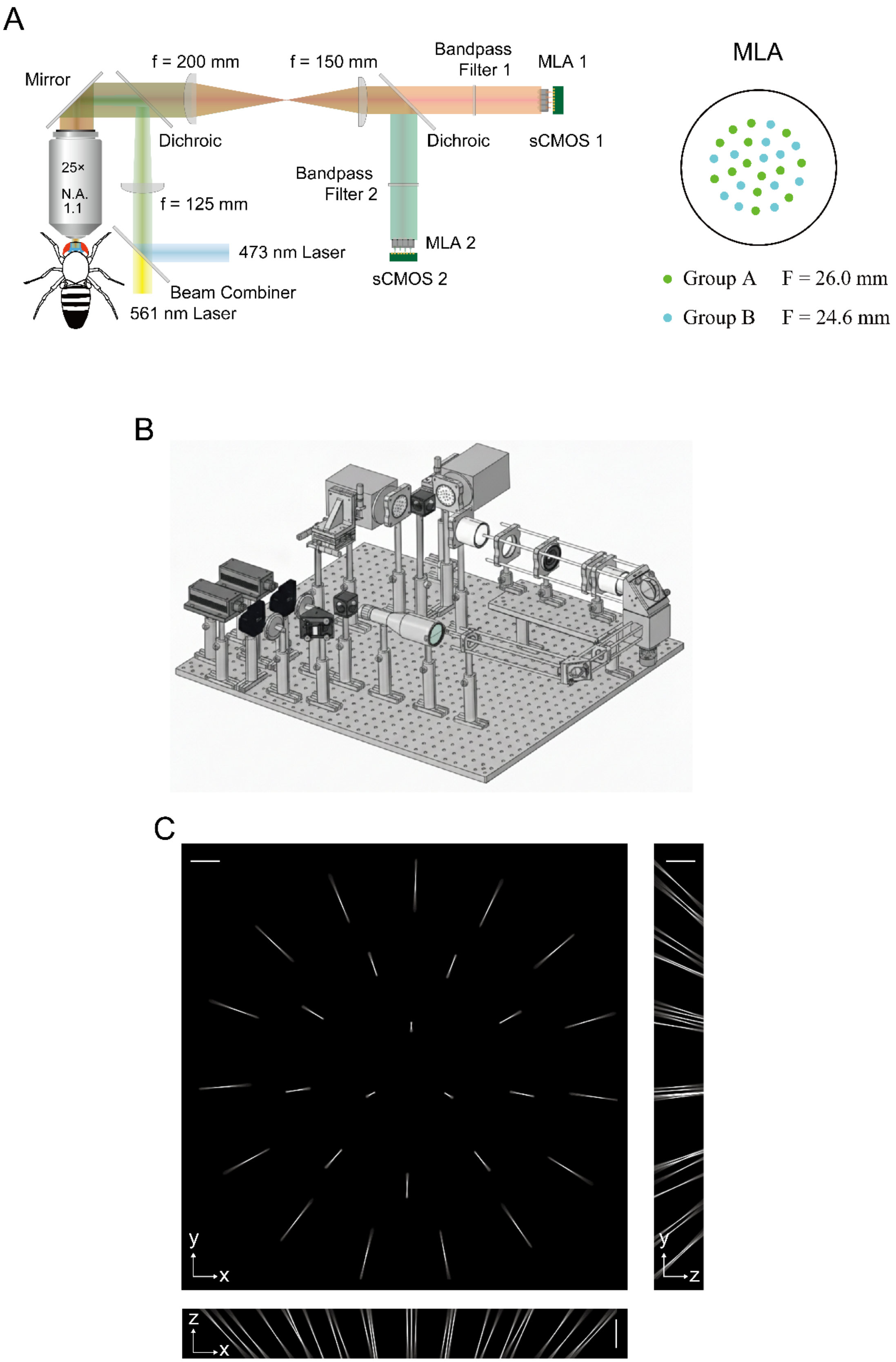
A
Mirror
f = 200 mm
f = 150 mm
Bandpass Filter 1
MLA 1
MLA
Dichroic
Dichroic
sCMOS 1
25×
N.A. 1.1
f = 125 mm
Bandpass Filter 2
473 nm Laser
MLA 2
Beam Combiner
561 nm Laser
sCMOS 2
Group A F = 26.0 mm
Group B F = 24.6 mm
B
C
y
x
y
z
z
x

**Figure S1 System design of dual-channel dual-focal Fourier lightfield microscope. Related to Figure 1.**

(A) Diagram of the optical system and the microlens array (MLA), with key parameters.

(B) Three-dimensional rendering of the optical system implementation.

(C) Measured PSFs of the system in the red channel, showing the maximum intensity projection of the PSF onto the x–y, y–z, and x–z planes. Scale bars are 200 μm. See Figure 1C for the PSF of the green channel.

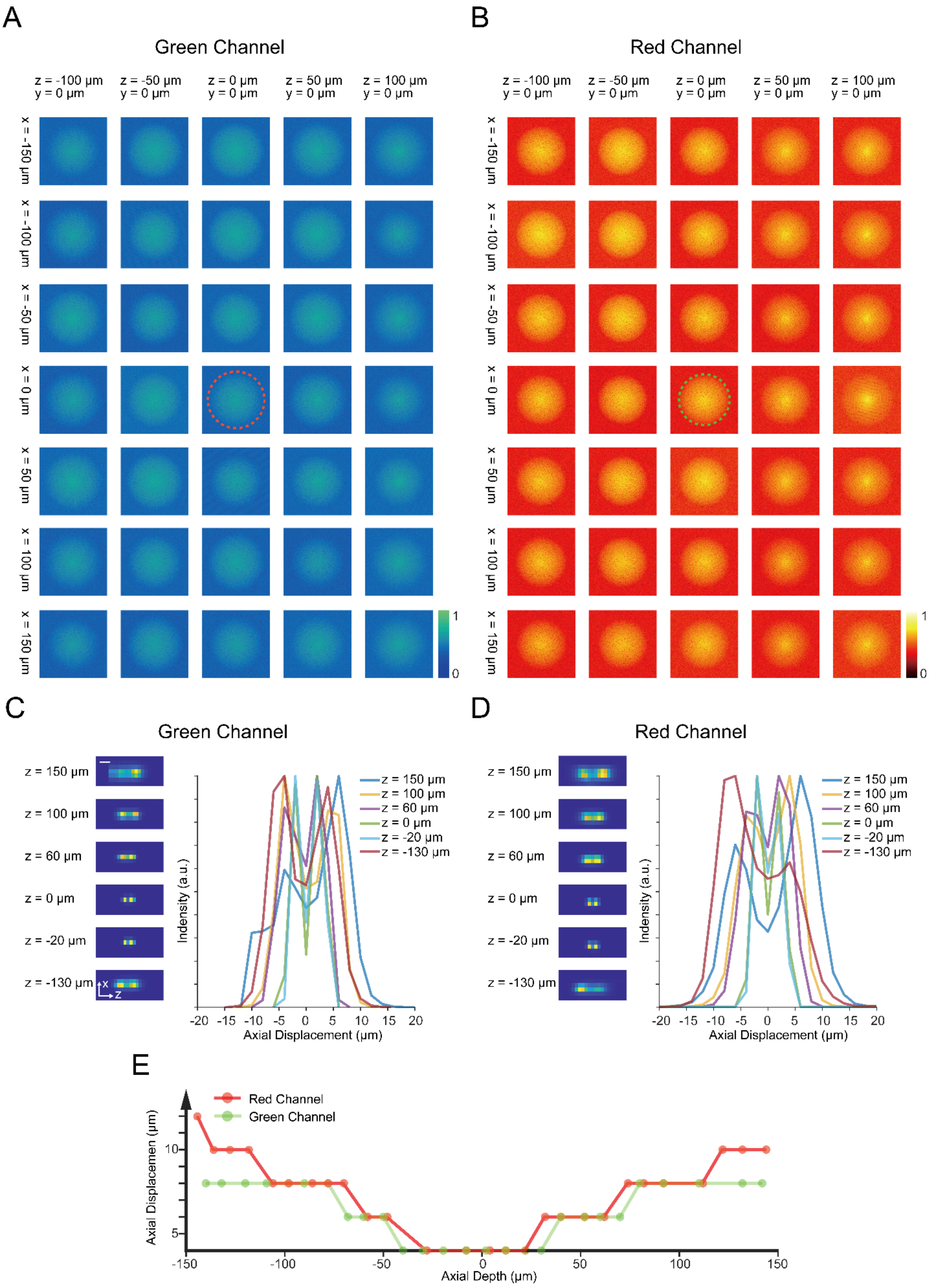

A
Green Channel
z = -100 µm y = 0 µm
z = -50 µm y = 0 µm
z = 0 µm y = 0 µm
z = 50 µm y = 0 µm
z = 100 µm y = 0 µm
x = -150 µm
x = -100 µm
x = -50 µm
x = 0 µm
x = 50 µm
x = 100 µm
x = 150 µm
B
Red Channel
C
Green Channel
z = 150 µm
z = 100 µm
z = 60 µm
z = 0 µm
z = -20 µm
z = -130 µm
Indensity (a.u.)
Axial Displacement (µm)
D
Red Channel
E
Red Channel
Green Channel
Axial Displacemen (µm)
Axial Depth (µm)

**Figure S2. Characterization of the spatial resolution of the dcFLF system. Related to Figure 1.**

(A-B) Characterization of micro-lens lateral resolution in the green (A) and red (B) channels. Log-scale Fourier transforms of raw images of 2-μm-diameter fluorescent beads. The particle was imaged at varying lateral (x = ±150, ±100, ±50, 0 μm) and axial (z = ±100, ±50, 100 μm) locations to evaluate the resolution of the micro-lenses across the field of view. Red and green dashed circles highlight spatial frequency components corresponding to the spatial resolutions.

(C-D) Characterization of the axial resolution in the green (C) and red (D) channels. To characterize the axial resolution at specific axial depths, the fluorescent bead was imaged at a series of axial locations with one particular axial displacement (3-15 μm) around the specific depth. The dcFLF volume was then acquired by synthesizing two raw images of a fluorescent bead with specific axial displacements around a given depth, followed by reconstruction. Each image on the left shows the X-Z projection of the 3D reconstructed dcFLF volume at particular depths (150, 100, 60, 0, -20, -130 μm). Curves on the right show the intensity profiles across axial displacements for each specific depth. Curves are color-coded by the axial depths. Scale bar is 5 μm.

(E) Axial resolution of the dcFLF system using the method in (C-D). The axial resolution at specific depths is defined as the minimal axial displacement required to resolve two particles distinctively.

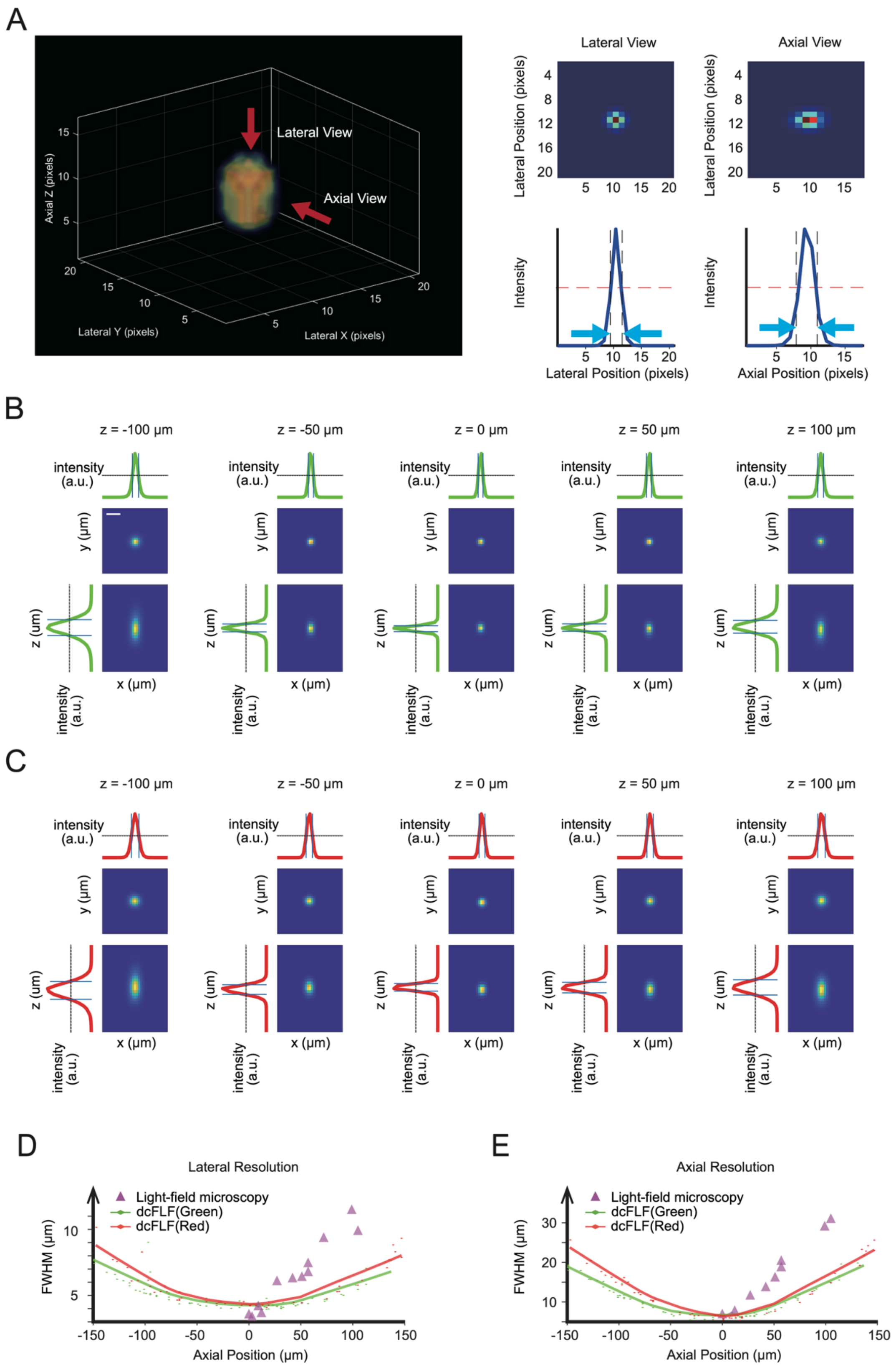
A
Lateral View
Axial View
Axial Z (pixels)
Lateral Y (pixels)
Lateral X (pixels)
Lateral Position (pixels)
Intensity
Axial Position (pixels)
B
z = -100 μm
z = -50 μm
z = 0 μm
z = 50 μm
z = 100 μm
intensity (a.u.)
y (μm)
z (μm)
x (μm)
C
D
Lateral Resolution
Light-field microscopy
dcFLF(Green)
dcFLF(Red)
FWHM (μm)
Axial Position (μm)
E
Axial Resolution

**Figure S3. Characterization of the spatial resolution of the dcFLF system via reconstruction. Related to Figure 1.**

(A) Illustration of the spatial resolution quantification. Left: 3D reconstruction of an example fluorescent bead, with fluorescence intensity color-coded. Top right panels: 2D projections of the 3D reconstruction onto two orthogonal planes. Bottom right panels: lateral (left) and axial (right) intensity profiles based on the lateral and axial view 2D projections. FWHM (blue dashes) was calculated from the intensity profiles to determine the spatial resolution.

(B-C) 2D projections of the 3D reconstruction onto lateral (top) and axial (bottom) planes at different axial depths in the green (B) and red (C) channels. Scale bar is 10 μm.

(D-E) Characterization of the lateral (D) and axial (E) resolutions of the system using reconstruction of imaged fluorescent beads. The horizontal axes denote axial positions relative to the native focal plane. The vertical axes show the FWHM of the intensity profiles as measurement of spatial resolution in the lateral and axial directions. Green and red diamonds and fitted curves denote the spatial resolution of the dcFLF in the green and red channels, respectively. For comparison, purple triangles indicate the spatial resolution of the conventional light-field microscope.

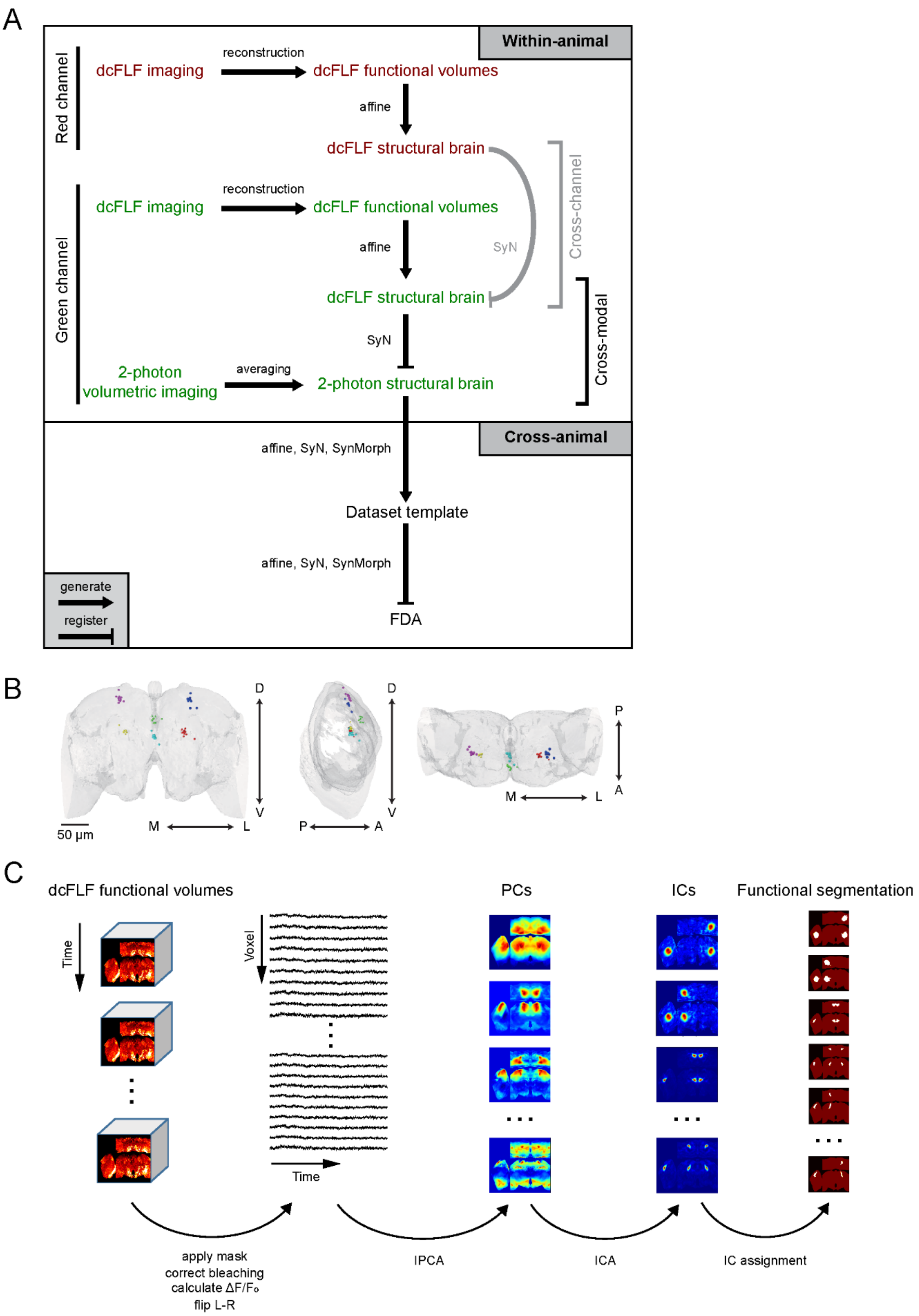

A
Within-animal
Red channel
dcFLF imaging
reconstruction
dcFLF functional volumes
affine
dcFLF structural brain
Green channel
dcFLF imaging
reconstruction
dcFLF functional volumes
affine
SyN
dcFLF structural brain
Cross-channel
SyN
2-photon volumetric imaging
averaging
2-photon structural brain
Cross-modal
Cross-animal
affine, SyN, SynMorph
Dataset template
affine, SyN, SynMorph
FDA
generate
register
B
D
V
M
L
50 µm
P
A
C
dcFLF functional volumes
Time
Voxel
PCs
ICs
Functional segmentation
apply mask
correct bleaching
calculate ΔF/Fo
flip L-R
IPCA
ICA
IC assignment

**Figure S4. Registration and segmentation. Related to Figure 2.**

(A) Registration pipeline, including within-animal registration and registration of the dataset. For each animal, cross-channel (gray) and cross-modal (black) alignments are performed sequentially. To obtain a structural brain for motion correction in each channel, subsets of dcFLF volumes from the time series are aligned using an affine transformation and then averaged into the structural brain. For cross-channel alignment, motion-corrected volumes of the two channels are aligned using affine and SyN transformation. For cross-modal alignment, a two-photon structural brain is similarly obtained using an affine transformation and averaging. Motion-corrected cross-channel aligned dcFLF volumes are then aligned with the two-photon structural brain using affine and SyN transformation.

For registration across animals in the dataset, cross-animal registration and registration to the FDA are performed sequentially. To perform cross-animal registration, aligned two-photon volumes from different flies are registered using affine, SyN, and SynMorph transformations to obtain the dataset template. To register the aligned dataset template to the FDA, we again used affine, SyN, and SynMorph transformations.

(B) Frontal, side, and top views of manually-labeled landmarks after registration. Each dot denotes the centroid of a labeled structure from one animal. The labeled structures include right (yellow) and left (red) BU, center between left and right mushroom body horizontal lobes (cyan), center of EB (green), top of right (purple) and left (blue) mushroom body vertical lobes.

(C) Pipeline of functional segmentation. Registered 3D volumes are subject to a series of preprocessing steps, including masking non-brain regions, bleach correction, and $\Delta F/F$ calculation. For data augmentation, the two hemispheres of each fly are flipped and concatenated with the original volume by the registered voxel. The 3D volumes of each fly thus obtained are then registered voxel-by-voxel into a single large-scale volume containing the augmented and combined dataset across flies. Incremental principal component analysis (IPCA) is then applied to this volume. This is then followed by independent component analysis (ICA) applied to the top principal components to extract independent components. Each voxel is then assigned the index of the independent component with the maximum loading as its cluster label. Each resulting cluster denotes a functional segmentation.

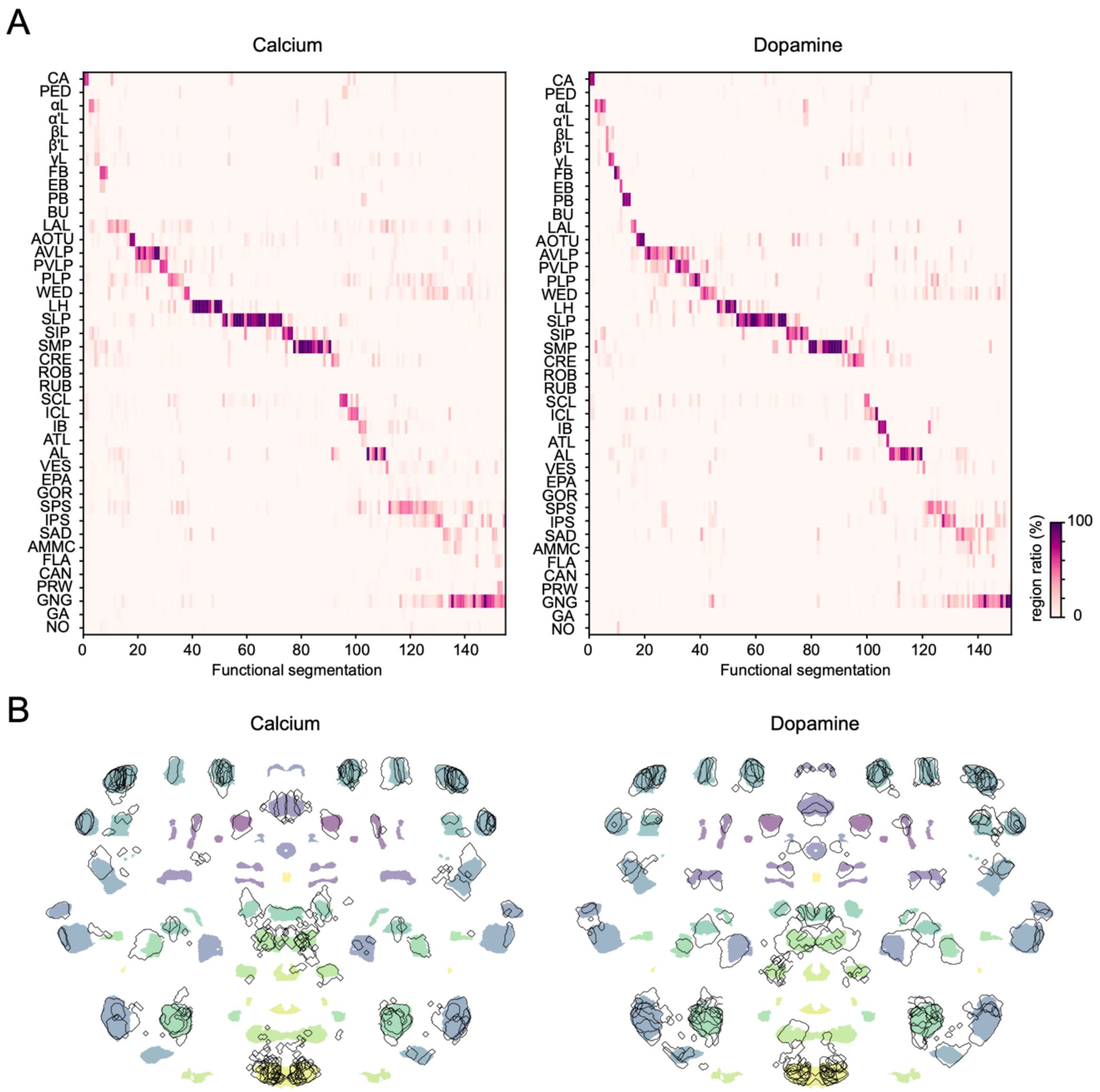
A
Calcium
Dopamine
CA
PED
αL
α'L
βL
β'L
γL
FB
EB
PB
BU
LAL
AOTU
AVLP
PVLP
PLP
WED
LH
SLP
SIP
SMP
CRE
ROB
RUB
SCL
ICL
IB
ATL
AL
VES
EPA
GOR
SPS
IPS
SAD
AMMC
FLA
CAN
PRW
GNG
GA
NO
0
20
40
60
80
100
120
140
Functional segmentation
region ratio (%)
100
0
B
Calcium
Dopamine

**Figure S5. Comparison between functional segmentations and anatomical regions. Related to Figure 3.**

(A) Relationship between anatomical regions (rows) and functional segmentations (columns) in calcium (left) and dopamine (right) channels. Color intensity shows the distribution of each functional segmentation across anatomical regions. Only segmentations with volumes larger than 0.7% of the brain are shown. See Table S1 for the abbreviations of brain regions.

(B) Morphological comparison between anatomical regions (colored shades) and functional segmentations (black outlines) in calcium (left) and dopamine (right) channels. Each functional segmentation is aligned with the anatomical region that shows the highest overlap. The brain is partially exploded. Segmentations and regions are visualized via volumetric projections onto the anterior-posterior axis. Only functional segmentations in which the anatomical regions with the highest overlap account for over 40% are shown.

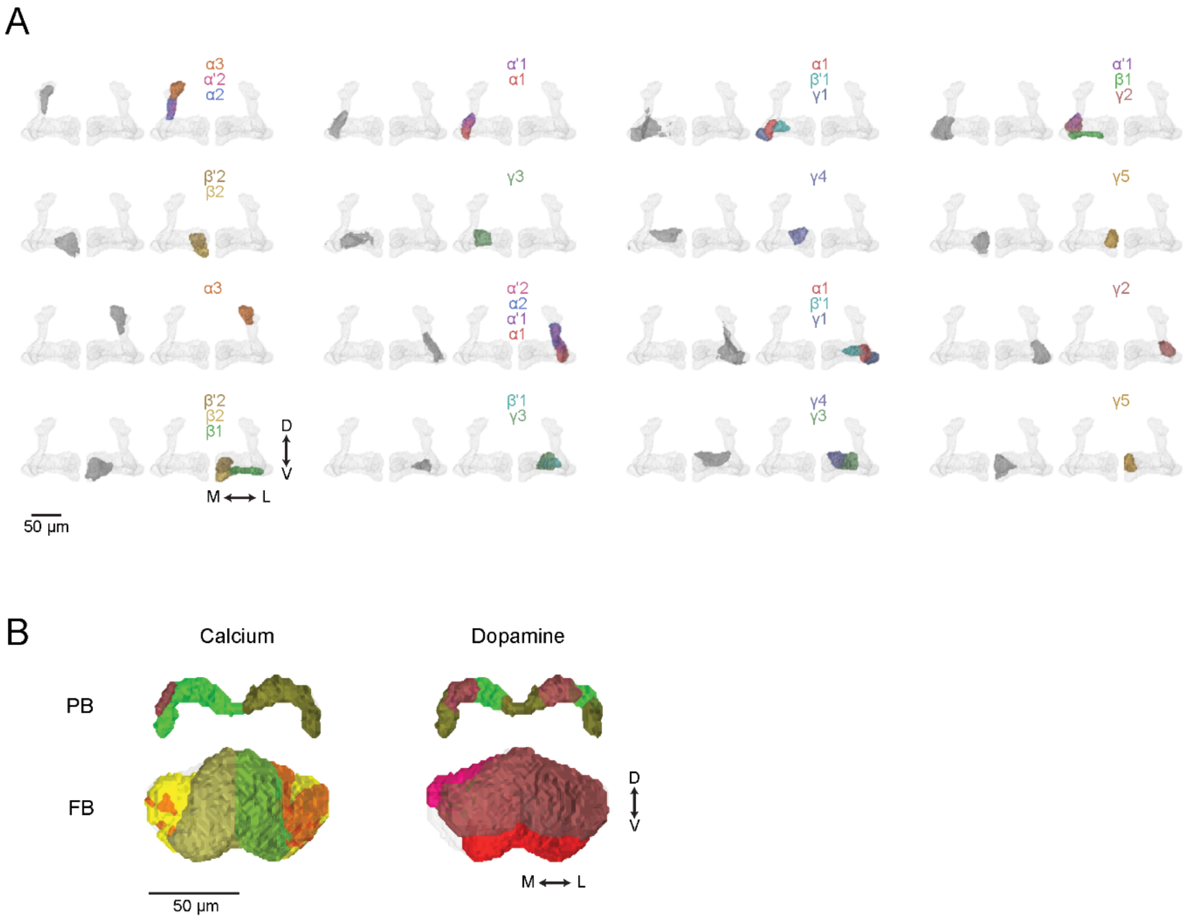

A
α3
α'2
α2
α'1
α1
α1
β'1
γ1
α'1
β1
γ2
β'2
β2
γ3
γ4
γ5
α3
α'2
α2
α'1
α1
α1
β'1
γ1
γ2
β'2
β2
β1
β'1
γ3
γ4
γ3
γ5
D
V
M
L
50 µm
B
Calcium
Dopamine
PB
FB
D
V
M
L
50 µm

**Figure S6. Functional segmentations in the mushroom body and the central complex. Related to Figure 3.**

(A) Comparison of 3D structures between anatomical compartments and functional segmentations in the dopamine channel in the mushroom body. For each functional segmentation (left, gray regions), the combination of compartments with the highest structural similarity is displayed and annotated (right, color-coded regions). Only segmentations larger than 5% of the mushroom body volume are shown.

(B) Comparison of 3D functional segmentations of the protocerebral bridge (PB, upper) and fan-shaped body (FB, lower) between calcium (left) and dopamine (right) channels. Colors denote functional segmentations. Note that FB exhibits a columnar segmentation for calcium but a layered segmentation for dopamine, while PB exhibits a left-right segmentation for calcium but multiple compartments for dopamine.

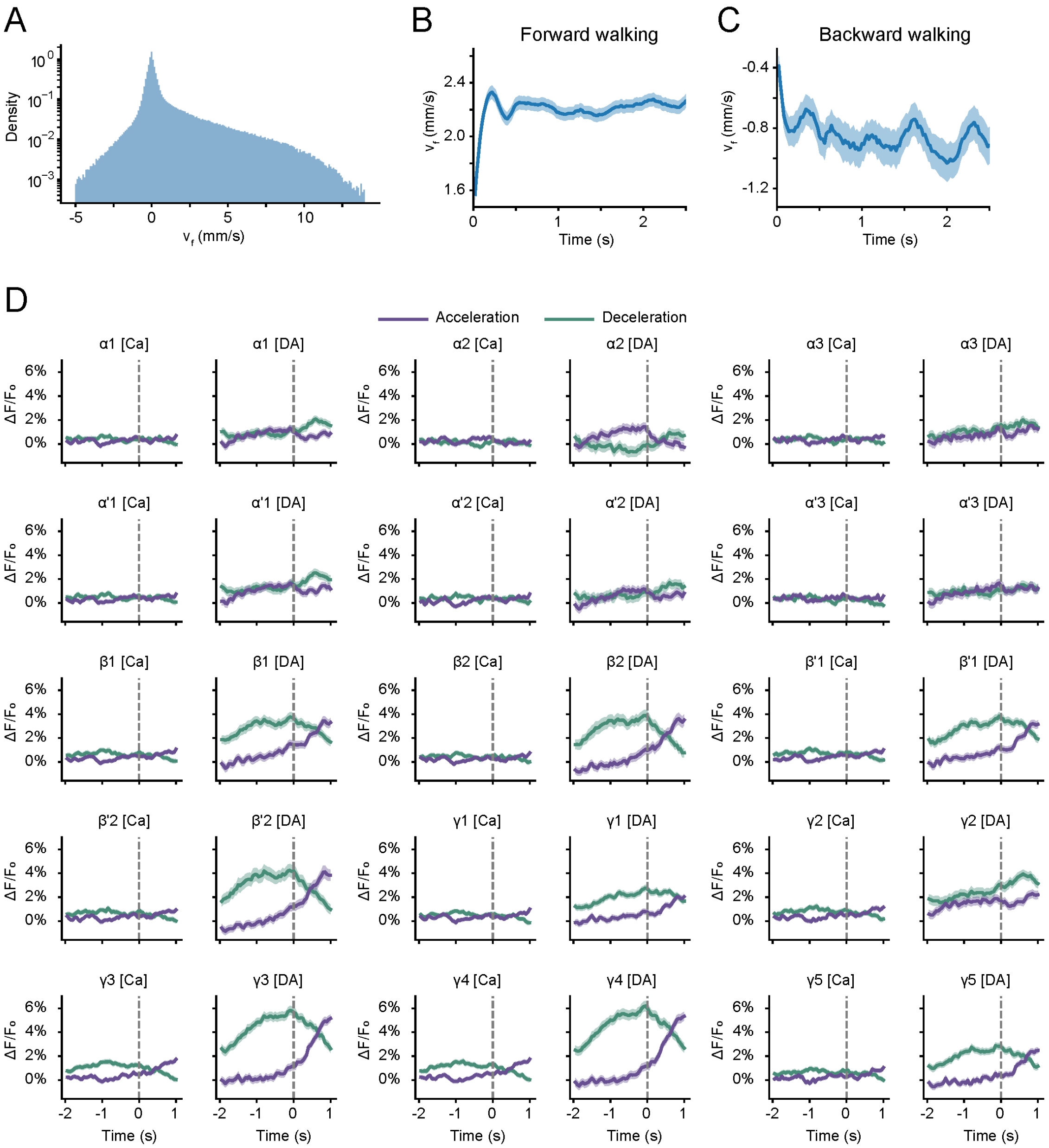

A
Density
v_f (mm/s)
B
Forward walking
v_f (mm/s)
Time (s)
C
Backward walking
v_f (mm/s)
Time (s)
D
Acceleration
Deceleration
α1 [Ca]
α1 [DA]
α2 [Ca]
α2 [DA]
α3 [Ca]
α3 [DA]
α'1 [Ca]
α'1 [DA]
α'2 [Ca]
α'2 [DA]
α'3 [Ca]
α'3 [DA]
β1 [Ca]
β1 [DA]
β2 [Ca]
β2 [DA]
β'1 [Ca]
β'1 [DA]
β'2 [Ca]
β'2 [DA]
γ1 [Ca]
γ1 [DA]
γ2 [Ca]
γ2 [DA]
γ3 [Ca]
γ3 [DA]
γ4 [Ca]
γ4 [DA]
γ5 [Ca]
γ5 [DA]
ΔF/F₀
Time (s)

**Figure S7. Calcium and dopamine activities in the mushroom body compartments during locomotion. Related to Figure 4.**

(A) Histogram of forward velocity distribution during the imaging experiments (n = 12 flies).

(B-C) Mean forward velocity dynamics of forward (B, n = 3477 trials from 12 flies) and backward (C, n = 173 trials from 12 flies) walking, shown as the mean ± S.E.M.

(D) Average calcium (Ca) and dopamine (DA) responses in mushroom body compartments during acceleration (purple) and deceleration (green) as in Figure 4F. Responses shown as the mean ± S.E.M.

| Abbreviations | Neuropil names |
|---|---|
| CA | calyx of mushroom body |
| PED | pedunculus of mushroom body |
| αL | mushroom body α-lobe |
| α'L | mushroom body α'-lobe |
| βL | mushroom body β-lobe |
| β'L | mushroom body β'-lobe |
| γL | mushroom body γ-lobe |
| FB | fan-shaped body |
| EB | ellipsoid body |
| PB | protocerebral bridge |
| BU | bulb |
| LAL | lateral accessory lobe |
| AOTU | anterior optic tubercle |
| AVLP | anterior ventrolateral protocerebrum |
| PVLP | posterior ventrolateral protocerebrum |
| PLP | posterior lateral protocerebrum |
| WED | wedge |
| LH | lateral horn |
| SLP | superior lateral protocerebrum |
| SIP | superior intermediate protocerebrum |
| SMP | superior medial protocerebrum |
| CRE | crepine |
| ROB | round body |
| RUB | rubus |
| SCL | superior clamp |
| ICL | inferior clamp |
| IB | inferior bridge |
| ATL | antler |
| AL | antennal lobe |
| VES | vest |
| EPA | epaulette |
| GOR | gorget |
| SPS | superior posterior slope |
| IPS | inferior posterior slope |
| SAD | saddle |
| AMMC | antennal mechanosensory and motor center |
| FLA | flange |
| CAN | cantle |
| PRW | prow |
| GNG | gnathal ganglion |
| GA | gall |
| NO | nodulus |

**Table S1. Abbreviations of neuropils. Related to Figure 3.**

Also see Figure S5A.
